# R2TP-like Quaternary Chaperones: a comprehensive overview to understand the dynamic R2SP complex

**DOI:** 10.1101/2025.01.27.635100

**Authors:** Paulo E. Santo, Marie-Eve Chagot, Hugo Gizardin-Fredon, Marie Ley, Thomas Chenuel, Evolène Deslignière, Laura Plassart, Ana C. F. Paiva, Pedro M. F. Sousa, Edouard Bertrand, Bruno Charpentier, Céline Verheggen, Marc Quinternet, Philippe Meyer, Tiago M. Bandeiras, Sarah Cianférani, Célia Plisson-Chastang, Xavier Manival

## Abstract

The human R2SP complex belongs to the R2TP-like quaternary chaperone family and consists of RUVBL1, RUVBL2, SPAG1 and PIH1D2. R2SP is crucial for the correct assembly of motile cilia (SPAG1 null mutations cause Primary Ciliary Dyskinesia) and the organization of the synaptic zone. RUVBL1/2 ATPases are the powerhouse of this molecular machinery, while SPAG1 and PIH1D2 would be adaptors that interact with specific clients to promote their quaternary assembly. Despite these functional data, little is known about the structure of R2SP and the precise mode of action of these R2TP-like complexes. We have combined biochemical and structural approaches (NMR, structural mass spectrometry and cryo-EM) to investigate the 3D organization of the human R2SP complex, its mode of assembly and ATPase activity. Our study reveals a three-dimensional structure similar to that of the canonical R2TP complex, but also highlights differences in the mode of action of its RUVBL1/2 ATPase core as well as the binding of its adaptors SPAG1 and PIH1D2.

## INTRODUCTION

Most proteins across all domains of life function as part of multimeric complexes, yet little is known about how their components assemble. In 2018, the term PAQosome (particle for <u>a</u>rrangement of <u>q</u>uaternary structure) was proposed to describe two *bona fide* multiprotein chaperones in mammalian cells - the R2TP and the Prefoldin-like (PFDL) complexes - that act together forming a megacomplex implicated in the assembly of quaternary structures crucial for cell growth^1–3^. The PFDL module includes PFDN2, PFDN6, PDRG1, UXT, URI1, ASDURF, the WD repeat-containing protein Monad (WDR92) and the RNA polymerase subunit POLR2E^4,5^. The R2TP complex cooperates with the HSP90 chaperone and includes the AAA+ (<u>A</u>TPases <u>a</u>ssociated with diverse cellular <u>a</u>ctivities) ATPases RUVBL1 (TIP49 or Pontin, R1) and RUVBL2 (TIP48 or Reptin, R2), RPAP3 (<u>R</u>NA <u>p</u>olymerase II-<u>a</u>ssociated <u>p</u>rotein 3, R3), and PIH1D1 (<u>P</u>rotein Interacting with <u>H</u>SP90 <u>1 d</u>omain-containing protein <u>1</u>, P1^6,7,8,2,9–11,3,12^). R1R2 are the catalytic core of the R2TP complex, while R3 and P1 are suggested to act as flexible scaffolding and regulatory units respectively. R2TP plays a role in the proper assembly of the growing number of protein or RiboNucleoProtein (RNP) complexes such as phosphatidylinositol 3 kinase-related kinases (PIKKs), RNA Polymerases, the C/D and H/ACA small nucleolar RNP particles (snoRNPs) or the U4 and U5 spliceosomal small nuclear RNAs. The existence of an endogenous R2T complex composed of R1, R2 and devoid of P1 has been shown both in human (owing to R3 isoform 2, that does not contain P1 binding site)^7,13^ and very recently in *Arabidopsis thaliana,* whose genome does not include P1^14^.

We identified a variant of the human canonical R2TP complex termed R2SP, responsible for the quaternary assembly clients involved in cell adhesion, cell migration, and organization of the synaptic zone^15^. The R2SP complex is composed by the R1R2 bound to two new protein adaptors, SPAG1 (<u>sp</u>erm-associated <u>a</u>nti<u>g</u>en <u>1</u>, S1), and PIH1D2 (<u>P</u>rotein Interacting with <u>H</u>SP90 <u>1 d</u>omain-containing protein <u>2</u>, P2). Paralogous to R3, S1 (104 kDa) has a larger N-terminal region, a third additional tetratrico<u>p</u>eptide repeat (TPR) domain and <u>R</u>3-Cter-like <u>b</u>inding <u>d</u>omain (RBD)^15,16^. 3D structural analyses show that TPR1 and TPR2 in R3 correspond respectively to TPR1 and TPR3 in S1, leaving an orphan TPR2^17,18^. The 100-residue linker region between the PBR and RBD in R3 is reduced to 25 residues in S1 (Fig. 1). P2 encompasses the same architecture as P1, namely a N-Ter phospho-motif domain which binds DpSDD/E consensus sites, and a C-terminal CS (<u>C</u>HORD-containing proteins and <u>S</u>GT1) domain required for S1 recruitment.

**Figure 1:**
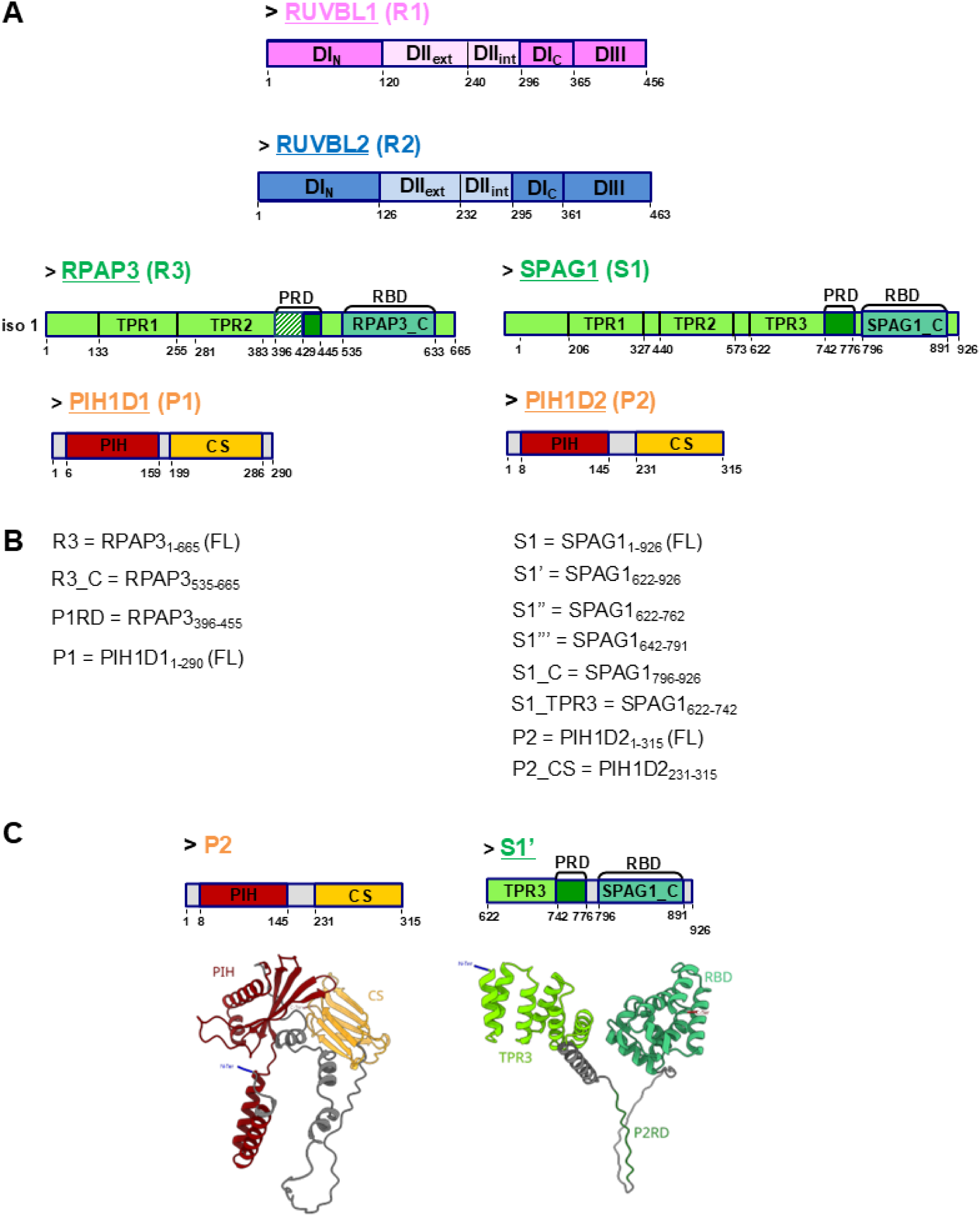
R2SP and R2TP complexes. **(A)** Domain organization of R2TP and R2SP components: R1, R2, R3 (in hatched, the insertion corresponding to isoform 1), S1, P1 and P2 proteins. **(B)** Sequence numbers of all protein constructs used in this work and their acronyms. **(C)** AlphaFold3 predictions of P2 (left panel) and S1’ (right panel) proteins.

R1R2 are the core engine within the R2TP and possibly R2SP, representing their generalist molecular machine-like role. Their atomic structure is preserved through species^16^ and presents an equilibrium between hetero-hexameric and hetero-dodecameric forms depending on nucleotide load and the nature of their partners. Each protomer comprises three domains. Domain I (D_I_) and domain III (D_III_) form the ATPase-core ring of R1R2 responsible for the ring-like oligomerization. A specific feature of R1R2 is the insertion of the domain II (D_II_) within the D_I_ primary structure. These flexible D_II_ protrude from the ATPase ring and are able to; (*i*) recruit cofactors/clients via their external region (D_IIext_) such as the DPCD (<u>d</u>eleted in a mouse model of <u>p</u>rimary <u>c</u>iliary <u>d</u>yskinesia) protein, which we recently identified^13,19–21^; (*ii*) establish a structural and conformational link via their internal region (D_IIint_) between clients and the ATPase core^20,22^; (*iii*) interact with nucleotides through their D_IIext_ region, which organize into oligonucleotide-binding (OB) fold domains^23^; (*iv*) interact with D_II_’s from other hexameric rings, forming D_II_-stacked hexameric rings (dodecamers), which are thought to represent an inhibited/storage form of the complex^6^.

Human R1R2 hetero-hexamers bind and hydrolyze nucleotides, and very often contain ADP in the active site pocket even when nucleotides were not intentionally added during the purification process^19,24^. The N-terminal parts of R1R2 interact via two conserved histidines with both their own D_IIext_ domains and the aromatic base and sugar moiety of the nucleotide, blocking the exchange from ADP to ATP^20,22^. Consequently, the absence of a nucleotide in the active side pocket results in the N-termini becoming unstructured. Whether and how nucleotides influence the way R1R2 recruit their partners is still a matter of debate. While O. Llorca’s group proposed that the recruitment of cofactors such as P1, leads to the opening of the R1R2 active site with concomitant ADP loss^20,25^, W. Houry’s group observed that the presence of P1 did not change the affinity of nucleotides for the R1R2 complex^13^.

R2SP/R2TP are both HSP70/90 (HSPs) co-chaperones specialized in quaternary protein folding during the assembly of key cellular macrocomplexes^1,15^. TPR1 from R3/S1 preferentially recruit HSP70 chaperones, and TPR2 from R3 and TPR3 from S1 preferentially recruit HSP90 by interacting with the common I/MEEVD motif located at the C-terminal tail of each chaperone^7,17,18^.

Depending on their specific adaptors, R2TP/R2SP target different protein or protein/RNA clients. The replacement of R3 and P1 in R2TP by the respective homologous proteins S1 and P2 in the recently discovered R2SP has a significant impact on the specificity of the resulting particle. Indeed, the R2SP complex seems to plays a role in the biogenesis of motile cilia via the quaternary assembly of axonemal dynein arms complexes^26–28^, while R2TP does not seem to be involved in this process.

3D structures of human^20,29^ and yeast^30,31^ R2TP, and plant^14^ R2T complexes have been recently published depicting the spatial arrangement of this eukaryotic canonical quaternary chaperone. The lack of a high-resolution structure for the full-length R2TP complex can be attributed to its high degree of flexibility and to the dynamic nature of its assembly. Here, we used biochemical, biophysical and structural approaches to characterize the human R2SP complex. Our work reveals similarities as well as significant differences compared to canonical R2TP, and hints towards its specific mode of assembly and action.

## RESULTS

### SPAG1:PIH1D2 synergistic effect on R2SP assembly differs from R2TP

To investigate its 3D structure and function, we have expressed all components of the human R2SP complex, separately or together, and used truncated versions when full-length (FL) proteins proved insoluble (Fig. 1). The C-terminal amino-acid sequences of R3 and S1 share 25% identity and 35% similarity scores, suggesting a conserved 3D fold for this domain and thus, putative conserved functions^17^. Based on the R3 C-terminal domain (R3_C) NMR structure^15^, we designed both a highly soluble fragment of S1 comprising residues 796 to 926 (S1_C), and a longer C-terminal soluble fragment from residues 622 to 926 encompassing the TPR3 and S1_C domains, which we named S1’ (Fig. 1)^17^. In our hands, FL SPAG1 (S1) can be stably over-expressed and was shown to bind co-chaperones HSP70 and 90 *in vitro*, however its ability to bind R1R2 proteins alone or in complex with P2 was not yet described^17^.

We first investigated the interactions between the different components of the R2SP complex *in vitro* using co-expression and co-immunoprecipitation assays and compared it with R2TP. The R1R2 complex co-purifies with all S1 construct variants, namely S1_C, S1’, and S1 (Fig. 2, A, B and C; lanes D, L and T) indicating that the S1_C is sufficient to bind the R1R2 complex, similarly to R3_C on R3 (Fig. 2, A and B; lanes H and P; controls are shown in fig. S1A). Additionally, NMR assays demonstrate that the presence of R1R2 degrades the ^13^C-METHYL-SOFAST spectrum of S1_C, regardless of nucleotide presence (Apo, ADP, or ATP) in the R1R2 active site pockets (fig. S2), indicating an interaction between S1 and R1R2 independent of nucleotide binding.

**Figure 2:**
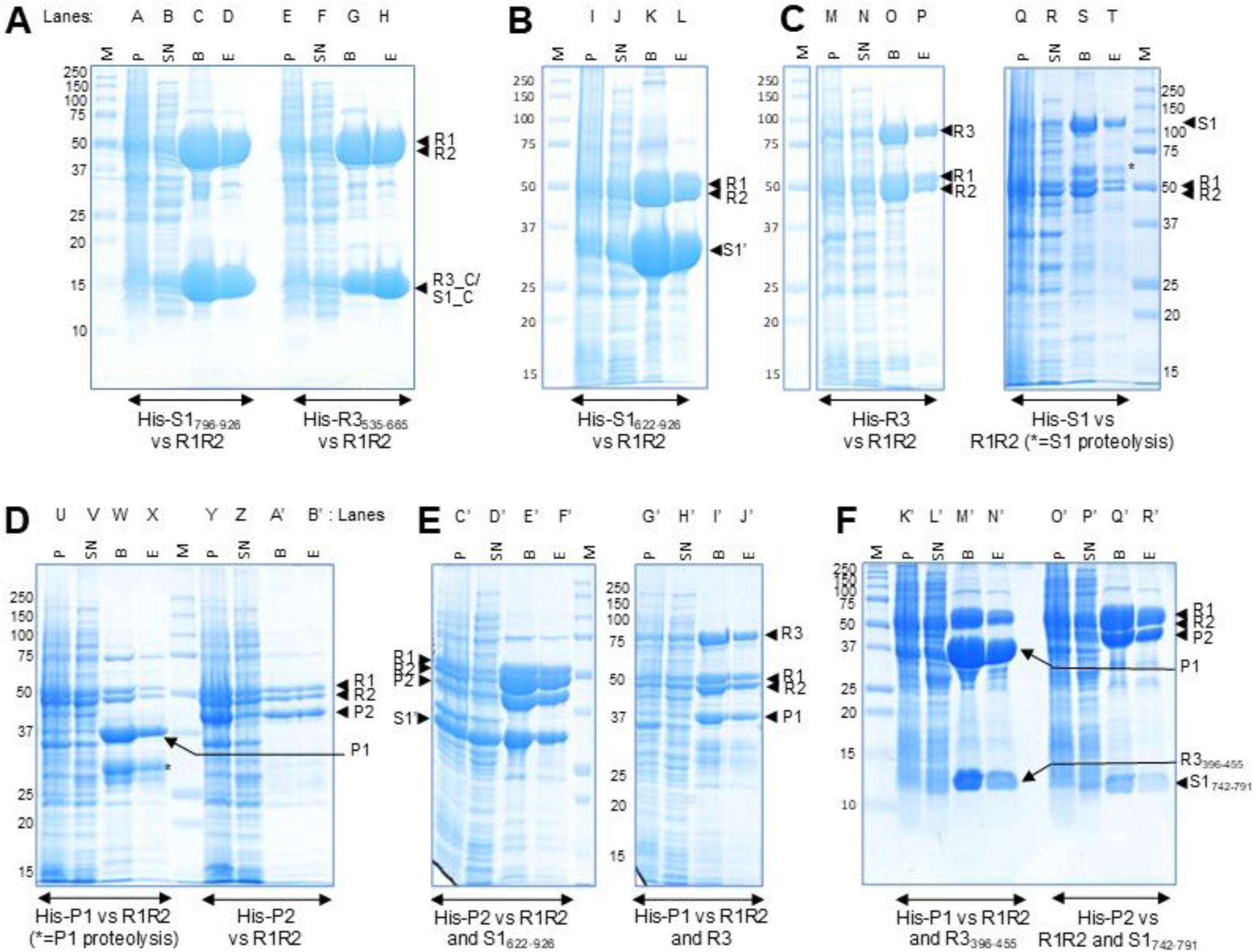
R2SP and R2TP co-expression assays. Coomassie blue stained SDS-PAGE of affinity purification on TALON resin of co-expression culture. Insoluble fraction (P), soluble fraction (SN), beads with proteins bound (F), elution (E) and molecular weight ladder (M) are loaded. **(A)** Co-expression of His-S1_796-926_ (His-S1_C) (lane A to D) and His-R3_535-665_ (His-R3_C) (lane E to H) with R1R2. **(B)** Co-expression of His-S1_622-926_ (His-S1’) with R1R2 (lane I to L). **(C)** Co-expression of His-R3 (lane M to P) and His-S1 (lane Q to T) with R1R2. **(D)** Co-expression of His-P1 (lane U to X) and His-P2 (lane Y to B’) with R1R2. **(E)** Co-expression of His-P2 with R1R2 and S1’ (lane C’ to F’) and His-P1 with R1R2 and R3 (lane G’ to J’). **(F)** Co-expression of His-P1 with R1R2 and R3_396-455_ (lane K’ to N’) and His-P2 with R1R2 and S1_742-796_ (lane O’ to R’).

As for both P1 and P2 proteins, their moderate co-elutions with R1R2 (Fig. 2D, lanes X and B’) suggest a lower affinity than R3 or S1 for the R1R2 heterocomplex. Note that P2 is mostly insoluble when co-expressed with R1R2 (Fig. 2D, lane Y), however unlike P1 that undergoes significant amount of proteolysis, the remaining soluble P2 (found in the supernatant and elution fractions) appears to be intact (Fig. 2D, lane B’ and X). P1 and P2 interactions with R1R2 become stronger with the additions of R3 and S1 to R2TP and R2SP complexes, respectively, with a more pronounced effect for R2SP (Fig. 2E, lanes F’ and J’). This suggests that P2 might first bind S1 for its efficient integration to the R2SP complex. Finally, we show that the S1 fragment 742-791 (downstream of the S1_TPR3 domain) can also integrate the R2SP complex, possibly through the CS domain of P2, similar to the binding of R3_396-455_ to P1^7^ (Fig. 2F, lanes N’ and R’). To further assess the interaction affinities within the R2SP complex, we performed Surface Plasmon Resonance (SPR) assays between different S1P2 sub-complexes, and different RUVBL variants namely: R1R2 complex (ADP loaded), double active site mutant R1^D302N^R2^D299N^ complex (no hydrolysis, ATP loaded), and R1R2 complex with truncated D_II_ (ADP loaded). The experiments demonstrate that the R1R2 complex has nanomolar affinity for both S1’P2 (K_D_ = 5.3 nM) and S1’P2_231-315_ (K_D_ = 22 nM) with resolved kinetics that allowed the determination of *k_on_* and *k_off_* (Fig. 3 and table S1). The R1R2 active site mutant loaded with ATP exhibits a kinetic profile for S1’P2_231-315_ which is similar to the interaction observed with the wild type, but once P2 is present in its FL form, the interaction profile changes, and the K_D_ increases to the micromolar range for the mutant form (fig. S3, A and C). The presence of ATP in the R1R2 active site has been related with D_II_ bending towards the active site (closed conformation)^20,22^, and this conformational change might stereochemically disfavor or occlude key regions for the interaction between the R1R2 mutant complex and FL P2, but not P2_231-315_, its CS domain. A similar effect is observed with the R1R2 dodecameric complex formed when D_IIext_ is truncated. In this case, either by the absence of D_II_ or by the occlusion of P2 interaction regions, low-affinity transient interactions are measured in the presence of P2 (fig. S3, B and D and table S1). Altogether, these interaction profiles suggest that D_II_ have a key role in the binding mechanism of adaptors S1P2 to the catalytic engine of R2SP complex, modulated by the nucleotide molecule present in the active pocket.

**Figure 3:**
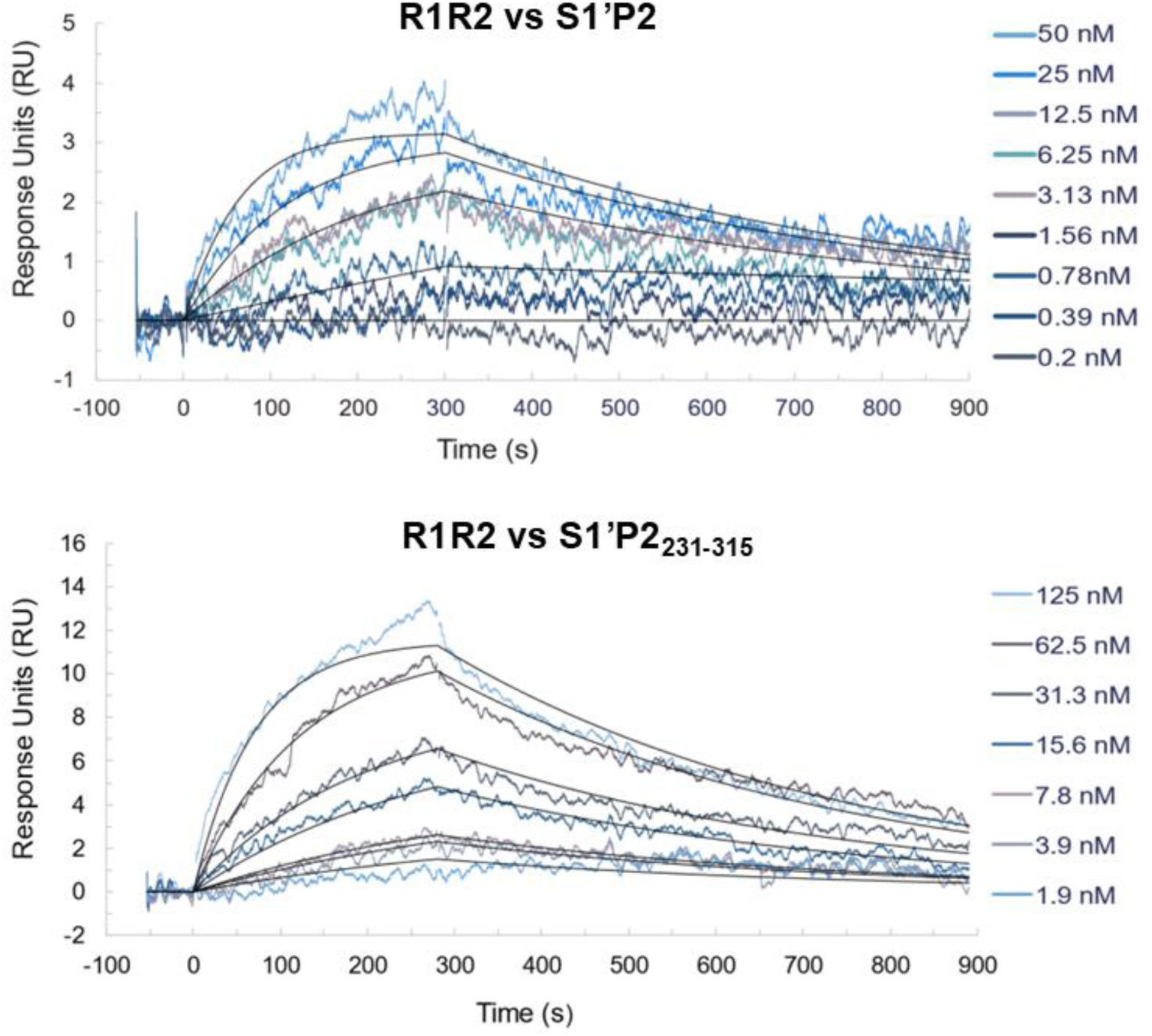
Experimental SPR sensorgram overlay plot from an injection series of R1R2 to a P2S1’ surface (Top) and P2_231-315_S1’ surface (Bottom) performed on a Biacore 4000. Colored curves represent recorded data and are overlaid with a 1:1 Langmuir binding model kinetic fit (black). The Top concentration and following injections are related by a 2-fold dilution. The K_D_ is taken as the ratio of the dissociation rate constant (*k*_d_) divided by the association rate constant (*k*_a_). Experiment was performed in absence of any nucleotide in the running buffer.

We further aimed to decompose the R2SP interactome and unveil its assembly mechanism by testing individual S1 or P2 proteins against the R1R2 complex. S1’ alone interacted very transiently with the R1R2 complex, with a determined K_Dss_ of 1.6 ± 0.2 µM (fig. S3, E and F), as previously shown by IP LUMIER assays *in vivo*^15^. Regrettably, S1, S1_C, and P2 failed to form functional active surfaces during the SPR assay development, even when using the Extract2Chip method^32^ which bypasses the hurdles of protein purification. Altogether, our data suggest that both S1 and R3 interact directly with the R1R2 complex^15^ but with different affinity (milli-vs nano-molar range, respectively). Moreover, S1 interaction with the R1R2 complex is stronger in the presence of P2, suggesting distinct cooperative models for the assembly of R2SP and R2TP.

### R2TP and R2SP exhibit different ATPase activity and ATP binding

We then compared the effect of S1P2 and R3P1 on the R1R2 ATPase activity. Using steady state kinetics, we demonstrate that both R3P1 and S1P2 can stimulate the R1R2 ATPases in a concentration-dependent and saturable manner, allowing the measurement of AC_50_ (concentration required for half-maximum activation) for both co-chaperone complexes (Fig. 4A). R3P1 can stimulate the R1R2 ATPase activity by 1.2-fold with an AC_50_ of 20 nM, in agreement with previous reports^13^, while S1P2 can stimulate R1R2 by 2-fold with an AC_50_ of 700 nM. These experiments suggest that although the S1P2 AC_50_ for R1R2 is higher than that of R3P1, S1P2 appears as a more potent activator of its ATPase machinery.

**Figure 4:**
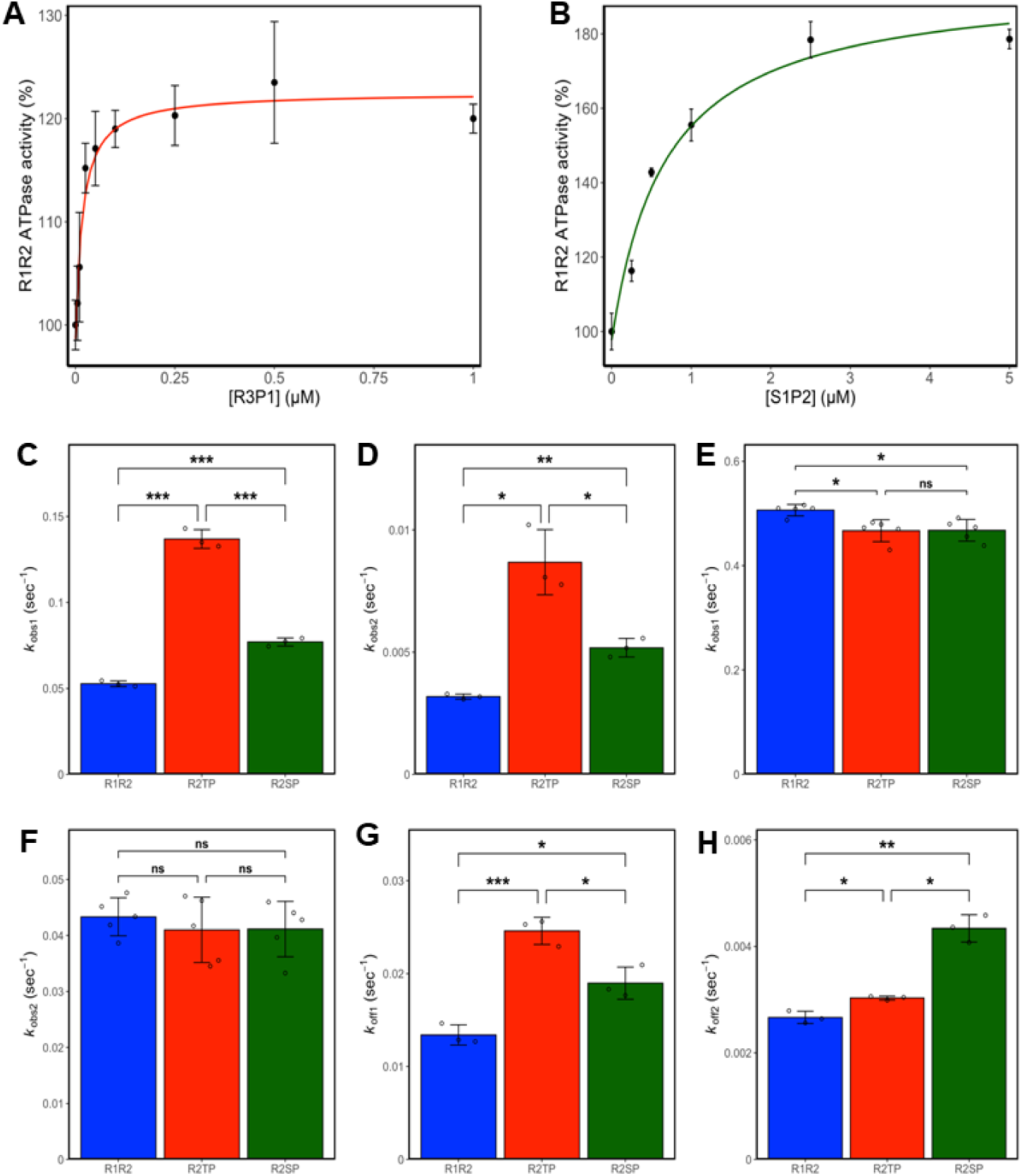
Dependence of R1R2 ATPase activity on R3P1 or S1P2 concentration. R3P1 **(A)** and S1P2 **(B)** concentration-dependent steady state ATPase activity of R1R2 was measured by a coupled assay. Data points (black dots) represent the means ± SD of three measurements at 37°C. The solid lines through data points represent the best rectangular hyperbola fit (rhombus for R3P1 and triangle for S1P2). **Kinetics of association and dissociation of fluorescent ATPγS on R1R2, R2TP & R2SP** was measured by a fluorescence polarization assay. Both showed a bi-exponential behavior (fig. S4). Apparent association constants *k*_obs1_ **(C)** and *k*_obs2_ **(D)** were extracted from best bi-exponential fits determined for R1R2 (blue), R2TP (red) and R2SP (green). A similar experiment was repeated on samples where bound ADP nucleotides were removed prior by alkaline phosphatase treatment and apparent association constants *k*_obs1_ **(E)** and *k*_obs2_ **(F)** with the same color code were determined. Finally, the nucleotide dissociation constants *k*_off1_ **(G)** and *k*_off2_ **(H)** were determined from best bi-exponential decay fits for R1R2 (blue), R2TP (red) and R2SP (green).

To decipher the mechanism of action of S1P2 and R3P1 on the R1R2 ATPases, we looked at their effect on nucleotide binding on R1R2. Using BODIPY-ATPγS (BDP-ATP), a fluorescent-labeled non-hydrolysable ATP analog, and fluorescent polarization measurement, we followed its binding and dissociation kinetics from R1R2 in the presence or absence of both co-chaperone complexes. We observed that nucleotide binding on R1R2 followed a bi-exponential regime suggesting that the R1R2 hexamer possesses two nucleotide binding sites with distinct association and dissociation rates, both differing by about 10-fold in magnitude and likely corresponding to the active sites of R1 and R2 (fig. S4). We named the two apparent association rate constants, *k*_obs1_ and *k*_obs2,_ and the two dissociation rate constants, *k*_off1_ and *k*_off2_, respectively. To characterize the effect of two co-chaperones on nucleotide binding, we first evaluated nucleotide association after the addition of either R3P1 or S1P2 (Fig. 4, C and D). We observed that both apparent association rates increase upon the addition of the co-chaperones, however R3P1 was 2 to 3-times more efficient in accelerating the association rates than S1P2.

Because our recombinant R1R2 preparation is associated with endogenous *E. coli* nucleotides (mostly ADP), we then checked if the *k*_obs_ would change if we first depleted nucleotides from R1R2. Indeed, we observed that the apparent binding rates on R1R2 alone are greatly accelerated by the absence of previously bound nucleotide (Fig. 4, E and F). Interestingly, the co-chaperones have no effect on the apparent binding rates but do accelerate nucleotide release (Fig. 4, G and H). Together, these observations suggest that R3P1 and S1P2 are more likely to facilitate nucleotide dissociation rather than nucleotide association. Although our experiments do not allow us to pinpoint which RUVBL has the slow or the rapid dissociating site, we observe that R3P1 is more potent in increasing *k*_off1_ of the rapidly dissociating site, while S1P2 is a better activator of the slow dissociation site and increases *k*_off2_. These differences could be related to different binding mechanisms of either R3P1 and S1P2 to the R1R2 AAA+ ring, already observed by SPR (this work and^15^).

### The C-terminal domain of SPAG1 and RPAP3 display similar 3D folds

To get further insight into the R2SP 3D organization and structurally characterize the putative R1R2 binding domain (RBD) of S1, we solved the NMR structure of S1_C in solution and obtained a well-resolved ensemble for which statistics are described in table S2. Briefly, S1_C is composed of 8 α-helices that pack together to form a globular domain (Fig. 5A). When compared to the NMR structure of R3_C, S1_C displays a similar overall fold, even if an extension of the loop between helices α7-α8 that covers helices 4 and 5 was observed in S1_C (Fig. 5B). Moreover, a break at the end of helix α_4_ was observed in both R3 and S1 RBDs. This break seems to represent a strong signature of this 3D fold. However, further comparison between R3_C and S1_C showed a RMSD value of ∼ 2.0 Å for Cα atoms in secondary structures. This suggests that S1_C may not be able to perfectly fit the binding site imprinted by R3_C on R1R2, even though the positioning of interfacing residues in R3_C appeared conserved in S1_C. This could help explaining why the SPR measured affinities with individual S1’ which contains the S1_C, (fig. S3), and R3_C^15^ for the R1R2 complex are different.

**Figure 5:**
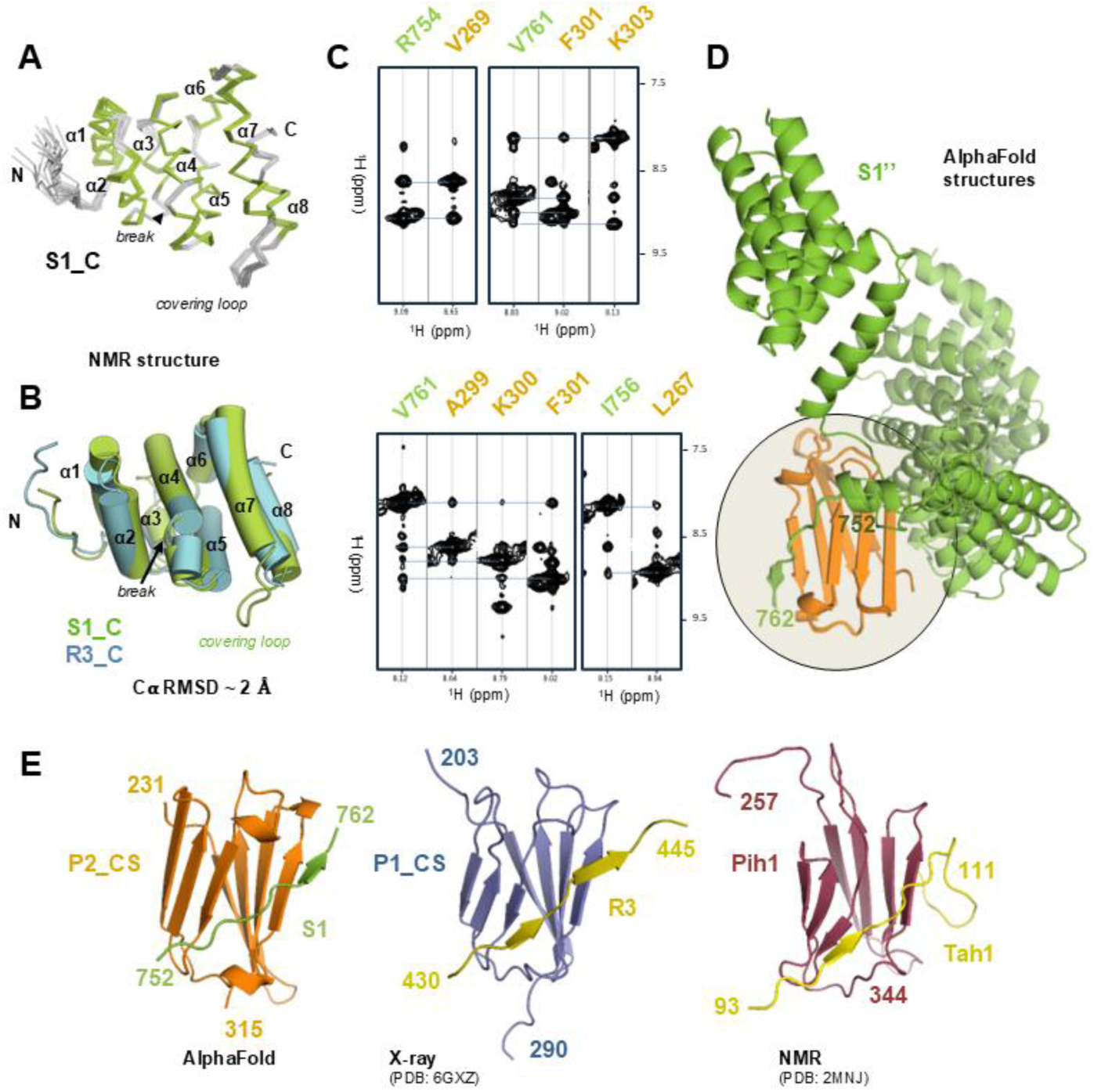
NMR structure of the C-terminal end of the human SPAG1 protein. **(A)** Backbone representation of the 20 best NMR structures (α-helices indicated in green). **(B)** Cartoon representation of the superimposition of the NMR structures of the C-terminal domain of human S1 (S1_C, green) and R3 (R3_C, blue) proteins. The break in the helix α4, common to S1 and R3, is indicated, as well as the covering loop, characteristic in S1. **(C)** Slices extracted from the ^1^H-^15^N NOESYHSQC spectrum recorded on a ^2^H/^13^C/^15^N-labelled P2_CS:S’’ complex. NOE cross peak linking amide protons belonging to individual spin systems are highlighted using dash blue lines. They reveal a spatial proximity between residues whose label is indicated on the top of the slice (green and orange label for S’’ and P2_CS, respectively). **(D)** Superimposition of the AlphaFold3 calculations of the P2_CS:S’’ complex. 3D structures were aligned on the 3D structure of P2_CS. The region of the complex with the highest convergence rate is highlighted in a circle. The fragment 752-762 of S1 is labelled. **(E)** 3D structures of the CS domain in P2, P1 and Pih1 in interaction with S1, R3 and Tah1, respectively. The structures come from AlphaFold3, X-ray and NMR data, respectively. The limits of each protein fragment are indicated.

### Validating the SPAG1:PIH1D2 interface using NMR and XL-MS constraints

Next, we aimed to characterize interactions between P2 and S1 at the atomic level. We successfully produced a ^2^H/^13^C/^15^N-labeled complex between the putative CS domain of P2 (P2_CS domain, P2_231-315_) and an extended version of S1_TPR3 (S1_622-762_, S1’’) (Fig. 1) and used NMR to perform the structural study. Despite the incomplete assignment of NMR peaks which prevents the full calculation of the complex 3D structure, experimental data showed that the 10-residue region 752-762 of S1 binds to P2 (Fig. 5C). The AlphaFold3 3D model of P2_231-315_:S1_622-762_ (P2_CS:S1’’) also reproduces this interaction between S1 and P2 (Fig. 5D) which are in very good agreement with intermolecular NMR distances and reveal the characteristic intermolecular β-sheet already observed in R3:P1 and Tah1:Pih1 (yeast ortholog) complexes (Fig. 5E). Interestingly, the positioning of S1_TPR3 on P2_CS is poorly conserved among the 3D structures returned by AlphaFold3, suggesting that these two domains do not strongly interact when expressed alone, or that they do so in a very dynamic manner. Our NMR and co-expression assays nonetheless led us to propose the 742-791 region of S1 as the minimal P2 binding region (Fig. 2F, lane R’).

In parallel, we performed XL-MS analysis of the full-length S1’P2 heterodimer using DSBU (DiSuccinimidyl diButyric Urea) as cross-linking agent, which can create bonds between lysine residues with Cα-Cα distances comprised between 6 and 32 Å (fig. S5). Overall, 58 unique cross-linked peptides were identified, distributed as 27 intra-S1’, 10 intra-P2 and 21 inter-S1’/P2 dipeptides (table S3 and fig. S5). Among those, 7 inter-S1’P2 dipeptides comprise the S1’_TPR3 domain involved in HSP70/90 binding and 11 inter-S1’P2 peptides are in the phospho-peptide binding pocket of the PIH domain (fig. S5).Interestingly, 1 inter-S1’P2 cross-links involve residues K_742_ from S1’ (close to its minimal P2 binding region) and K_293_ from P2 (in its CS domain), which complements the S1’P2 binding interface observed in NMR. downstream of the S1’_TPR3 domain, a region described to be involved in P2 interaction (Fig. 2F, lane R’).

### Up to three S1’:P2 heterodimers interact with one R1R2 heterohexamer

We used co-expression to assemble a native yet truncated version of the R2SP complex in bacteria (fig. S6). Because the full-length S1 protein suffers proteolysis, we used the S1’ construct (Fig. 1) and produced the R2S’P complex by mixing pre-assembled R1R2 with S1’P2, in the presence of ADP, and used a gel filtration as a final polishing step to purify the quaternary complex, as described previously for the R2TP complex^29^. To assess the stoichiometry of the R2S’P complex, we used two complementary biophysical techniques: native mass photometry (nMP) and native mass spectrometry (nMS). While nMP profiles obtained for R1R2 alone revealed the coexistence of hexameric and dodecameric assemblies (fig. S7), the R2S’P nMP histograms revealed a much more heterogeneous distribution, with a majority of hexamers compared to dodecamers (73% vs. 27% of oligomeric assemblies, respectively) (Fig. 6, A and B). Similarly, comparison of nMS spectra obtained for R1R2 and R2S’P (fig. S7B and Fig. 6C) also demonstrate that S1’P2 binding disrupts the R1R2 dodecamer to form a stable R2S’P complex based on hexameric R1R2, like binding of other R1R2 partners (e.g. DPCD or RPAP3^19,20^). For hexameric species, both approaches revealed the coexistence of undecorated R1R2 hexamers (313,503 ± 8 Da, 20%), or carrying one (384,692 ± 13 Da, 35%) or two (456,039 ± 22 Da, 18%) S1’P2 units, each containing an average of 6 ADP molecules (Fig. 6C). A fourth very low-abundance population (< 5%), that could correspond to a R1R2 hexamer with three S1’P2, can be seen but not accurately mass assigned. These stoichiometries are similar within the R2TP complex^20^ and highlight that the R1R2 can recruit several S1P2, although its implications *in vivo* are still poorly understood.

**Figure 6:**
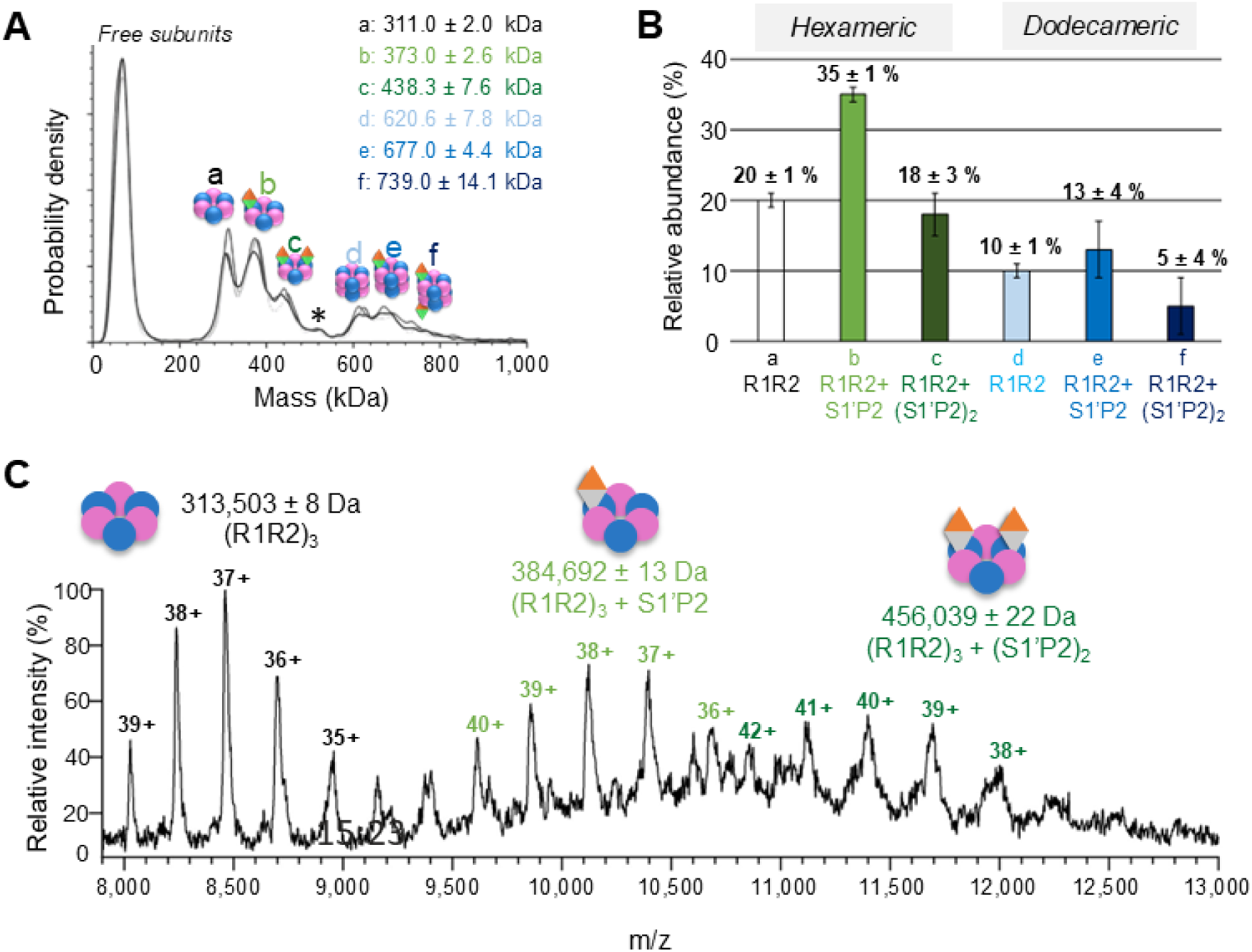
Determination of R2SP oligomeric states using Mass Photometry and native MS. **(A)** Mass Photometry measurements in triplicates, shown in shades of grey, at a droplet concentration of 50 nM. MP shows that R1R2 hexamers co-exist with R1R2:S’P2 and R1R2:(S’P2)_2_, and that dodecameric R1R2 is also formed. Standard deviations come from measurement replicates (n=3). In the 30-100 kDa mass range, free R1 or R2 monomers overlap with S1’P2 heterodimer. In the 100-550 kDa mass range, three populations are identified and quantified: undecorated R1R2 hexamers (311.0 ± 2.0 kDa, 20 ± 1%), or R1R2 with one (373.0 ± 2.6 kDa, 35 ±1%) and two (438.3 ± 7.6 kDa, 18 ± 3%) S1’P2 bound. Above 550 kDa, populations of R1R2 dodecamers (620.6 ± 7.8 kDa, 10 ± 1%) with one (677.0 ± 4.4 kDa, 13 ± 4%), or two (739.0 ± 14.1 kDa, 5 ± 4%) S1’P2 bound are dectected. * represents low abundance R1R2:(S’P2)_3_ species (<5%). **(B)** Relative abundances of the different oligomeric states, in % of total counts, measured in MP. Standard deviations come from measurements replicates (n=3). **(C)** native MS confirmed the presence of R2S’P complexes with different stoichiometries ranging from R1R2 hexamers bound to one (384,692 ± 13 Da) or two S’P2 (456,039 ± 22 Da), each containing in average 6 ADP molecules.

### R2S’P and R2TP complexes share similar 3D architecture

We next investigated the 3D structures of *in vitro* reconstituted human R2S’P using cryo-electron microscopy (cryo-EM) and single particle analysis (SPA). Initial 2D and 3D classification rounds using RELION^33^, revealed the coexistence of dodecameric and hexameric assemblies (fig. S8), in agreement with MS characterization. The image processing scheme allowed to reconstruct a 3.5 Å map of the dodecameric assembly (fig. S9). It is formed by two toe-to-toe R1R2 hexamers and appear skewed, as already observed by other groups for full-length R1R2 assemblies^19,29^. Conformational heterogeneity studies of the R1R2 dodecamer using cryoDRGN^34^ also shows movements of one hexamer relative to the other (movie S1), which likely reflect the structural dynamics between the two hexamers as previously suggested^35^. Of note, 2D classes as well as low-pass filtered cryo-EM maps of dodecameric assemblies reveal horn-like densities crowning the AAA+ core of R1R2 (fig. S10), which could correspond to the dynamic binding of S1’, as hinted by our nMP and nMS observations and further cryo-EM analyses.

After having sorted out particles, we further classified and refined the alignment of hexameric particles, and this processing scheme yielded a 3D reconstruction of R2S’P solved to 3.6 Å resolution (fig. S11). We used this cryo-EM map (Fig. 7A) and the NMR structure of the S1_C fragment described above to build an atomic model of the R1R2S1’ assembly (Fig. 7B). It revealed the presence of at least one “horn” on top of the R1R2’s D_I_-D_III_ hexameric ring that could be attributed to the RBD of S1’. Like for its paralog in R3, this domain straddles one R1 and one R2 monomers (Fig. 7, A and B). Note the presence of the three R1’s D_IIext_ in the open conformation, and the higher flexibility of the R2’s D_IIext_ where S1’ is bound on D_I_-D_III_ upper ring.

**Figure 7:**
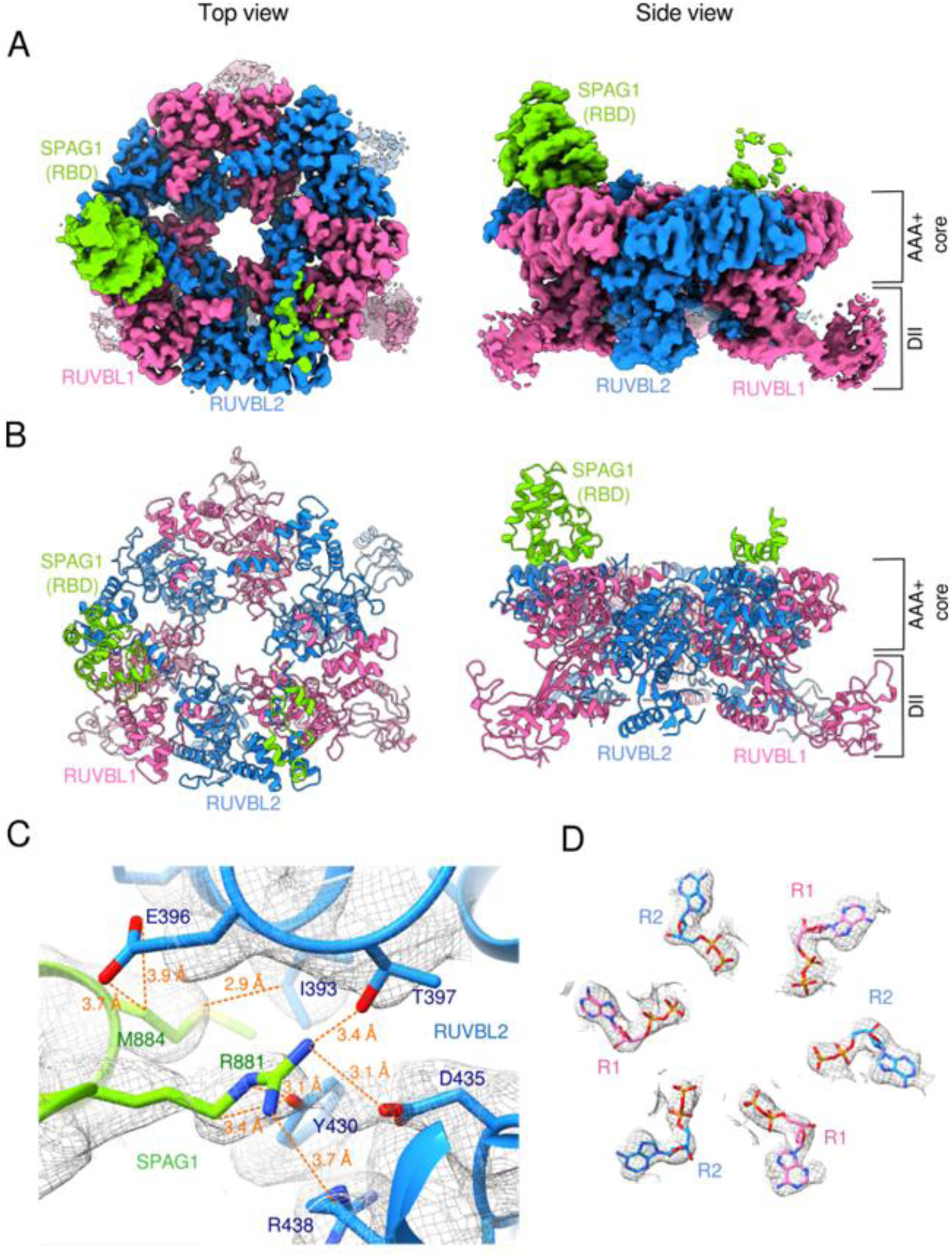
3D structure of the hexameric R2S’P complex. **(A)** Top (left panel) and side (right panel) surface views of the cryo-EM map of hexameric R1R2 solved at 3.6 Å resolution. **(B)** Corresponding views of the atomic model derived from the cryo-EM map shown in A. R1 and R2 are shown in pink and blue, respectively. S1_C is shown in lime green. **(C)** and **(D)** Details of the cryo-EM map (grey mesh) and reconstructed atomic model of the contact zone between S1_C and R2 **(C)** and of the ADP moieties in the R1R2 hexamer (D). Colour coding is the same as panels (A) and (B).

Inspection of the R2S’P cryo-EM map shows a close proximity (<4 Å) between S1’ residue R_881_ and R2’s R_392_, T_397_, Y_430_, D_435_ and R_438_, while M_884_ from S1’ establishes contacts with (at least) R2 residues E_396_ and I_393_ (Fig. 7C). Moreover, in agreement with nMS data, 6 ADP molecules could unambiguously be fitted into the cryo-EM density (Fig. 7D). Interestingly, a density on top of the D_I_-D_III_ domains of the R1R2 hexameric ring was attributed to partial presence of a second S1’ molecule (Fig. 7, A and B), in agreement with nMS results. To further analyze the stoichiometry of S1’ on R1R2 particles, we performed focused classification of the set of hexameric particles (fig. S11), which showed that the R1R2 heterohexamer could indeed be crowned by either 1 or 2 S1’ (fig. S12), with a particles proportion of 63% and 37%, respectively. In parallel, we used cryoDRGN^34^ to sort out the structural heterogeneity of the hexameric dataset using AI-based methods. This revealed R1R2 heterohexamers crowned by one, two or even three RBD domains of S1’ (movie S2, PC1 analysis), in agreement with nMP and nMS results.

Low-pass filtered 3D classes (fig. S12B) as well as 2D classes (fig. S12C) of hexameric R2S’P particles always reveal fuzzy densities (that were never refined to high resolution despite our many trials) below R1R2’s D_IIs_, which seem to be centered compared to the inner cavity below the R1R2 ring (fig. S12B and movie S2, PC1 analysis). These densities were also clearly detected by cryoDRGN analysis of the binned cryo-EM images (movie S2, PC1 analysis). By homology with the R2TP complex^20^, we propose these densities to correspond to the dynamic association of P2.

Based on these results, we next tested different surface interaction mutants suggested to be fundamental for the specific interaction between R2 and S1’, namely: D435 for R2 and R881A/M884A for S1. The S1’s mutations correspond to residues R_623_ and M_626_ in R3, which are essential for its interaction with R1R2^15^. We used both co-expression (fig. S13) and IP-LUMIER (fig. S14) assays to demonstrate that the aspartic acid 435 of R2 is essential for its interaction with S1 and R3. The co-expression results show that the interaction is conserved between R1R2^D435A^ mutant and R3_C or R3 (fig. S13, B and D). They also show that the addition of P2 restores the interaction between mutant forms of R2 and S1 up to the formation of R2SP complex, which is not completely the case with P1 (fig. S13, F and G, and S14B). This is in good agreement with our SPR results, in which the presence of P2 is also important for S1 to bind the R1R2 complex. Here again, our analysis confirms a key interplay between S1 and P2 upon binding to the R1R2 hexamer to form the R2SP complex, while the assembly determinants appear different for the R2TP complex. These differences might be core reasons explaining the R2TP and R2SP specificity *in vivo*.

### XL-MS reveals the importance of RUVBL1/2’s D_II_ domains in SPAG1 and PIH1D2 interaction

To complement cryo-EM data, we studied in-solution interactions formed within the R2S’P complex with XL-MS analysis, using DSBU as a cross-linker. A total of 208 unique cross-links (99 intra- and 109 inter-protein interactions) were identified in the R2S’P complex and visualized in a network plot (Fig. 8A and table S4). Direct interactions between P2 and S1’ with R1R2 were detected confirming R2S’P complex formation. Almost 37% (40 peptide pairs) of the unique inter-XLs involve P2 and either R1 (26) or R2 (14). Half of them interact via the D_II_ domains of R1R2, and within these, 14 involve the PIH domain of P2. The XL-MS data supports that P2 interact with R1R2 not only via their D_II_ domains, as suggested by interactions and SPR studies, but also via their PIH domains. On the S1’ side, the network plot also reveals many interactions between S1’ and R1R2 (27 cross-links pairs, 25% of the total inter-XL pairs), mostly involving the D_III_ domain of R2 (residues K_368_, K_417_ and K_427_) while all R1 domains (D_I_, D_II_ and D_III_) are engaged in R1-S1’ interaction (Fig. 8B, table S4). While observing several S1’ cross-links with the different domains of both RUVBLs, we propose that although firmly anchored via its S1_C to R2, the remaining S1’ construct is relatively flexible, which also helps explaining why it is hard to reconstruct in cryo-EM data. In addition, XL-MS dataset also revealed 17 inter S1’-P2 dipeptides, mainly between the CS domain of P2 (K_252_, K_293_ and K_306_) and the TPR3 of S1’ (K_641_, K_670_, K_706_ and K_710_), in agreement with NMR results obtained on smaller constructs. Finally, the PIH domain of P2 is also in close proximity of S1’_TPR3, as depicted by the cross-links between K_42_ of P2 and K_628_/K_744_ of S1’ (fig. S15).

**Figure 8:**
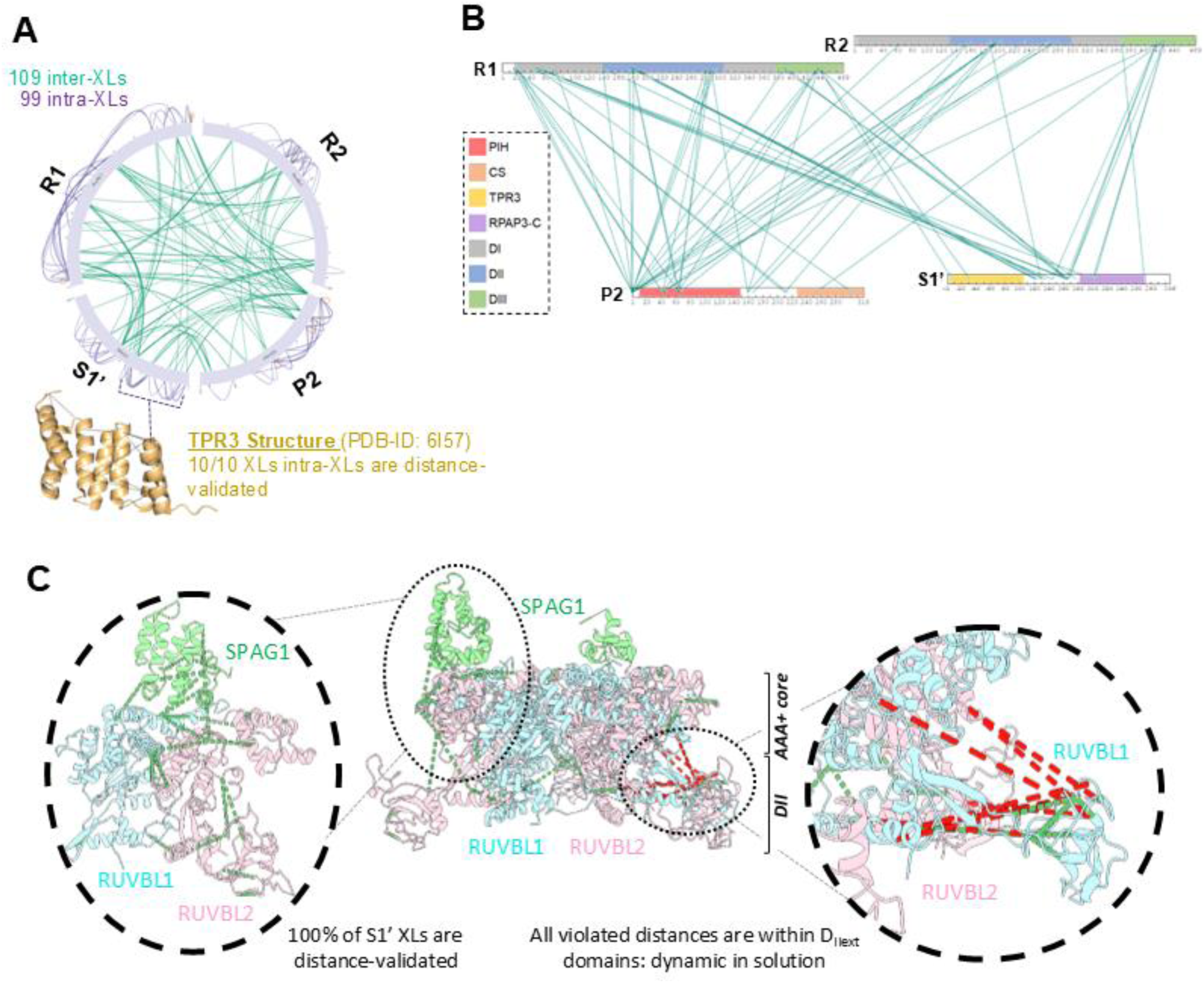
Description of R2S’P XL-MS. **(A)** Circular plot showing identified intra- and inter-XLs in R2S’P dataset. All intra-S1’ XLs of TPR3 domain could be mapped and distance-validated on NMR TPR3 S1’ structure (6I57.pdb). **(B)** Interaction network between R1, R2, S1’ and P2 in R2S’P as identified by XL-MS. **(C)** XL-mapping on R2S’P resolved cryo-EM structure, with XL distance-validated in green and violated in red. Overall, 84% of mapped XLs are in accordance with structure. The S1 positioning and conformation are confirmed by 4 inter-XLs and 4 intra-XLs, all distance-validated.

We next mapped the identified cross-links on the cryo-EM maps, (Fig. 8C). As the high-resolution cryo-EM reconstruction lacks P2 and is poorly resolved in RUVBLs D_IIext_ regions, only 51 XL pairs (24%) could be mapped on the EM structure (30 intra-links and 21 inter-links). Among these inter-XLs, 10 involve R1 and R2, while 5 account for S1’-R1R2 pairs. Remarkably, 84% of measurable Cα-Cα distances on the EM structure where in agreement with DSBU distance constraints (6-32 Å). The violated distances (8 XL-pairs) all include D_IIext_, already known to be dynamic in-solution (fig. S16). MS dataset thus cross-validate EM results and, especially S’1 positioning within the cryo-EM structure (Fig. 8C).

### An integrative 3D model of the R2S’P complex

We first performed an AlphaFold3 structure prediction of the R2S’P complex leading to a similar overall structure for R2S’P compared to the cryo-EM structure. However, the D_IIext_ of R1 and R2 are always found in a closed position in AlphaFold3 predictions (fig. S17A), contrarily to our cryo-EM data, where D_IIext_ seem to be moving along a plane that is perpendicular to the R1R2 central tunnel and not a parallel one (see movie_S2, PC2 analysis). Representation of our XL-MS data on the AlphaFold3 model led to 58% of XL-pairs distances validated, with no agreement with predicted inter S1’-P2 interface (11/12 violated) and less than a half (11/27) of S1’-R1R2 XLs distance-validated (fig. S17B), which questions the positioning of S1’s TPR and P2’s CS domains in this AlphaFold3 model.

Considering those discrepancies with our experimental data, we next combined NMR, cryo-EM and XL-MS data with AlphaFold3 structure predictions of P2 and/or S1’P2 complexes, to propose a 3D model of the R2S’P complex (Fig. 9, fig. S18). Positioning of S1’ was done using 7 XLs with R1 and 4 with R2, while P2 positioning was done using 5 XLs with R1 and 2 XLs with R2 (fig. S18). We could globally plot on the integrative model 148 XLs out of the the 208 XL-pairs identified in XL-MS dataset, corresponding to a 3-times increase compared to the cryo-EM structure of R2SP with missing densities (51/208). Out of these 148 XL-pairs, 71% were distance-validated, supporting overall positioning of proteins in integrative model (Fig. 9B). S1’ and P2 conformations were distance-validated with 100% and 64% of intra-protein XLs, respectively. Altogether, this integrative architectural model suggests that the S1_C and PIH domains of S1 and P2 would clamp the R1R2’s ring hexamer at their top and bottom sides, respectively, while S1’s TPR3 and P2’s CS domains would interact together in the vicinity of and with the D_IIext_ of R1R2 (Fig. 9A).

**Figure 9:**
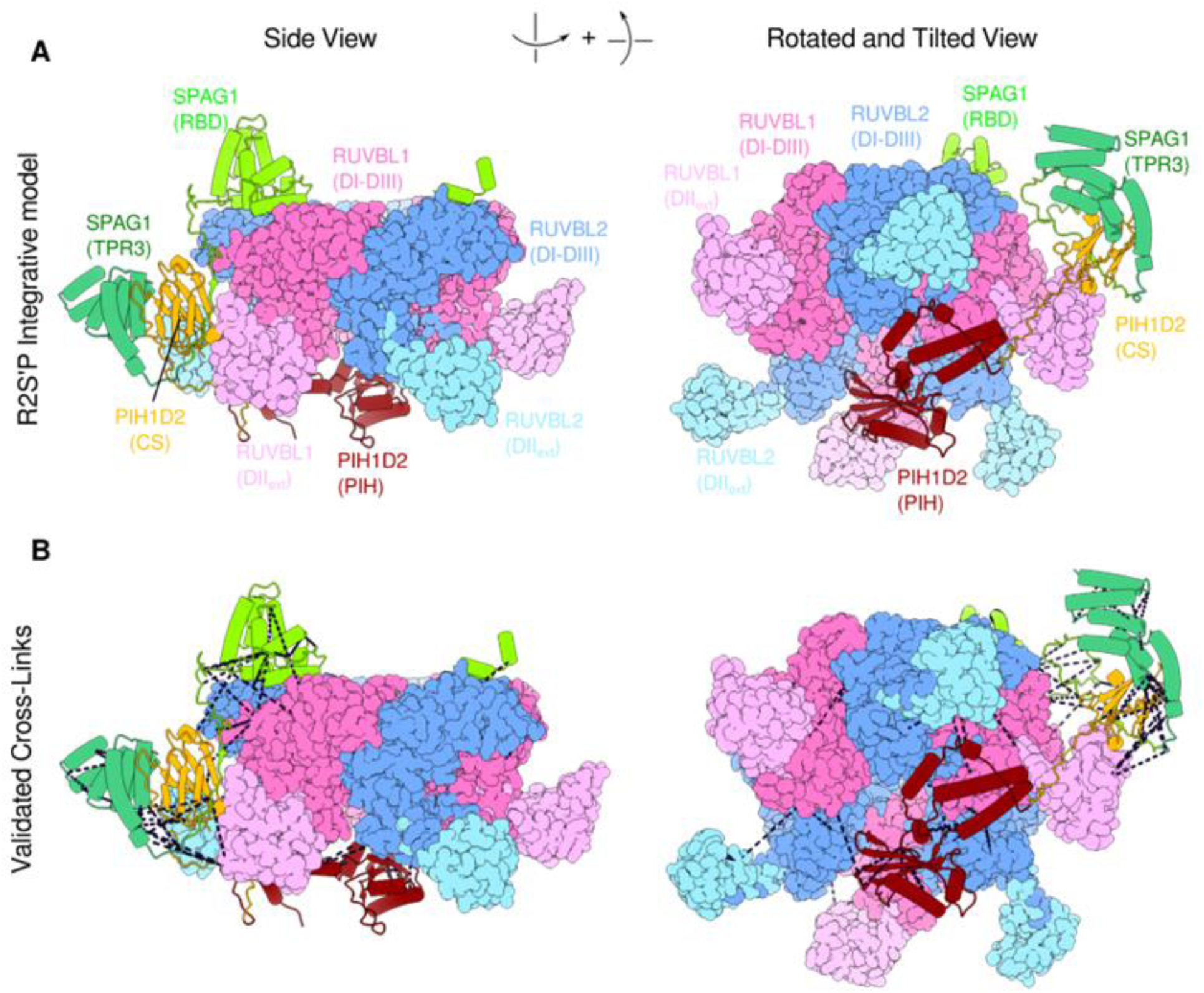
Integrative 3D model of the R2S’P complex. **(A)** hypothetical 3D model of the full R2S’P assembly based on NMR, XL-MS, cryo-EM and AF3 predictions. RUVBL1 and 2 are represented in pink and blue, respectively, the RBD and TPR3 domains of SPAG1 (S1’) are shown in lime and forest green, respectively. PIH1D2 is represented in dark red (PIH domain) and orange (CS domain). **(B)** Representation of cross-link pairs validated on the integrative model (71% of the total XL pairs, black dashed lines).

## DISCUSSION

In this study we used biochemical, biophysical and integrative structural biology approaches to unveil for the first time, mechanistic and architectural aspects of the quaternary chaperone R2SP, which has been shown to play a role in motile cilia biogenesis via the quaternary assembly of axonemal dynein arms^26–28^. The association of R1R2 with cilia components is most likely bridged by the already known cilia interactors S1 and P2^15,36,37^, while the molecular mechanisms of how this work remain elusive, as for all R1R2 complexes.

As S1 and P2 share several homologous domains with R3 and P1 proteins, which belong to the archetypal R2TP complex^7^, we herein performed a comparative biophysical study of human R2TP and R2SP. Within R2TP, R3 is the sole binder to the R1R2 complex^13,15^. W. Houry’s group further suggest that P1 does not directly interact with R1R2 in the presence of R3, but is rather positioned near the ring due to its direct interaction with R3^13^. This is further supported by the existence of sub-complexes such as R2T and R2P^13^ with yet unknown functions, and the existence of an endogenous version of R3 (isoform2) unable to interact with P1^7^. Such stable sub-complexes for R2SP and splicing isoforms for S1 have not yet been identified *in vivo*, and our data indicates that they might not form at all. Furthermore, our co-expression and SPR analysis hint towards a cooperative binding mechanism of S1 and P2 to the R1R2 ring, as S1 seems to require P2 to recreate a high affinity interaction. This is somehow reinforced by our point mutation experiments, which highlight how the same mutation in R2 only affects S1 binding, but not R3. Hence, our results pinpoint to a different assembly mechanism for the R2SP as compared to the R2TP, and we would however need to further elucidate P2’s role upon binding to the R1R2 complex (this was technically not achievable due to P2 alone being unstable in the SPR assays). Nevertheless, our data clearly indicate that the physico-chemical properties of P2 (from R2SP) differ from P1 (from R2TP), by playing a crucial role in the assembly of the R2SP complex.

ATP binding and/or hydrolysis are required for most of the R1R2-related quaternary chaperones^38^. Our activity results show that S1P2 stimulates the R1R2’s ATPase activity almost twice as much as R3P1. This might be correlated with our SPR analysis, in which the use of ATP-loaded dead mutant versions of R1R2 negatively impacts the interaction with the S1’P2 sub-complex when compared to WT, *i.e.* ADP-containing R1R2. Altogether, these data indicate that the ATP binding to the R1R2 active site, and S1P2 association to R1R2 are strongly interdependent, but how the ATP presence in the active site influences the S1’P2 binding kinetics or vice versa remains elusive. Does the nucleotide lead to changes in the D_I_ and D_III_ ring’s apical region, interfering with the S1 binding? Or does it influence D_IIext_ positioning as previously proposed^22^, in turn affecting P2 binding? Furthermore, our kinetic assays show that both R3P1 and S1P2 co-chaperones would facilitate nucleotide dissociation rather than association to R1R2 binding sites. Our experiments on R1R2 alone, further reveal the presence of two types of nucleotide binding sites, each with distinct association and dissociation rates (one faster than the other). R3P1 appears more potent in dissociating non-hydrolysable ATP from the fast site, while S1P2 is more efficient on the slower site. We hypothesize that those two ATP binding sites correspond to R1 and R2, but further experiments are needed to identify which one of the two is the slower or the faster. Of note, in our cryo-EM structure (which was built without imposing symmetry), all active sites are fully occupied by ADP molecules, regardless of being bound to S1 or P2, suggesting a different molecular mechanism from R2TP, where based on the previous solved cryo-EM structure^20^. It is suggested that P1 promotes nucleotide release from the active site of an R2 protomer, while the other ring’s active sites remain ADP bound. However, another group^13^ using ATPase assays showed no significant effect of R3P1 alone in the R1R2 rings activity.

The three-dimensional structure of the R2SP complex was totally unknown, and could only be inferred by comparison with its homologous R2TP complex^20^.Our cryo-EM, XL-MS and NMR experimental data allow to propose a three-dimensional model of the R2SP chaperone that closely resembles R2TP, in which P2’s PIH domain, known to recruit phosphorylated clients^39^, is located below the ring formed by the D_IIs_ of R1R2 (basal ring side). In contrast, its CS domain appears bound close to S1’s TPR3 domain, and both of these are located close-by the R1R2’s D_II_ external parts, most probably on the side of R1R2 hexameric assembly. This is favored by different lengths in the linker regions present in-between P1 and R3 structured domains, compared to P2 and S1 pair. Indeed, R3 has a long linker spanning from the motif that binds the CS domain of P1 (PRD) to its RBD (Fig. 1), which was previously proposed to play a role in the competition between R3 and P1 direct binding to the R1R2 D_IIext_^13^. In contrast, S1’s shortened PRD linker might impose the P2 protein to adopt a bilobal conformation with the CS domain near the R1R2 ring’s vicinity, while the PIH domain is in the basal region, which could help explain why the R2SP assembly mechanism seems to differ from the R2TP’s one. We hypothesize that the bilobal conformation adopted by P2 could compensate for the shorter S1 linker region, and thus facilitating S1 binding to R1R2.

Based on our R2SP 3D model, it can be proposed that client complexes assembled by the R2SP complex will be done so on the D_II_ side of the ring, recruited through P2. S1, which is anchored via its C-terminal RBD to the opposite side of the R1R2 ring would thus keep HSP70/90 chaperones (known to bind the N-terminal TPR domains located just before the PRD^17,18^) “hanging” close by to assist the assembly process. Of note, no clients have been shown to bind the upper side of the R1R2 ring (S1_RBD side) up to present day, and this binding polarity might be extended to other quaternary folding processes involving R1R2.

Finally, our nMP, nMS and cryo-EM results show that similarly to the R2TP^13,20^, several S1P2 decorate the R1R2 ring. Our extensive R2S’P cryo-EM image analysis also hints towards a high flexibility of R1R2 D_IIext_, P2 and S1’s TPR3. Altogether, we propose that this spatial organization allows R2SP to synchronize the interaction of its different clients, their transfers from HSP70/90 and that R2SP could be considered as an octopus-like dynamic platform for quaternary client assembly. Our study of the human R2SP complex adds another biophysical and structural brick to the characterization of the compendium of R1R2-based quaternary chaperones, shading light into chaperoning molecular mechanisms deeply dependent on fine-tuned protein:protein interactions.

## METHODS

### NMR

NMR protein samples were prepared with a classic procedure involving, (*i*) production in *E. coli* BL21(DE3) pRARE2 cells grown in a minimal M9 media containing ^13^C-D6-glucose and ^15^NH_4_Cl, and (*ii*) a two-steps purification process based on immobilized metal ion affinity chromatography followed by gel filtration. For the P2_CS:S1’’ complex, the bacterial cultures were performed in D_2_O.

A classic set of 3D NMR spectra was recorded at 298 K using a 600 MHz AVANCE III spectrometer equipped with a cryoprobe. This enabled the nearly complete ^1^H, ^13^C, and ^15^N resonance assignments for S1_C, but only a partial assignment of the backbone of P2_CS:S1’’. For structure calculation of S1_C, backbone chemical shifts were converted into dihedral angle restraints using TALOS-N. The CYANA 3.97 automated procedure was then employed to extract distance restraints from 2D ^1^H-^1^H NOESY, 3D ^15^N-NOESY-HSQC, and 3D ^13^C-NOESY-HSQC spectra, all recorded with a mixing time of 120 ms. The resultant sets of dihedral angles and inter-proton restraints underwent thorough verification and were utilized to generate 100 CYANA structures, subsequently refined in explicit water using the AMBER-based Portal Server for NMR structures (AMPS-NMR). The 20 structures with the lowest constraint energies were chosen as the most representative. Visualization of all 3D structures was performed using Pymol software^40^. Chemical shifts were deposited in the Biological Magnetic Resonance Bank under reference 34968. The 3D structure of S1_C was deposited in the Protein Data Bank under reference 9HKR.

To monitor the binding of unlabeled RUVBL proteins to ^13^C-labeled S1_C, 1D METHYL-SOFAST-HMQC spectra were recorded. Interaction experiments were conducted at 298 K and 600 MHz in the R1R2 buffer (20 mM Tris-HCl pH 8.0, 250 mM NaCl, 5% glycerol, 2 mM MgCl_2_, 0.5 mM TCEP). R1 was tagged with a 6xHistidine at the N-terminus, and R2 was tagged with a 3xFLAG. The concentration of ^13^C-labeled proteins was approximately 10 μM. The ^1^H dimension was edited to selectively detect ^1^H nuclei attached to ^13^C nuclei. Protons attached to ^13^C nuclei in the range of 5 to 35 ppm were selectively excited with a bandwidth of 3 ppm centered at 0 ppm. The relaxation delay was set to 150 ms, and the number of scans was 2048. The final concentration ratio between unlabeled and labeled proteins was maintained at 1:1, considering one monomer of S1_C and one heterodimer of R1R2.

### LUMIER IP

HEK293T cells were seeded in 24-well plates and transfected with 450 ng of the RL fusion and 50 ng of the 3xFLAG-FFL fusion, with 1 μL of JetPrime (PolyPlus), as recommended by the manufacturer. 48 h later, cells were extracted for 15 minutes at 4°C in 450 μl of HNTG containing protease inhibitor cocktail (Roche) and spun down at 4°C and at 20,000 x g for 10 minutes. The IP was performed duplicated, by putting 100 μL of the extract in each of four wells in a 96-well plate, with two wells being coated with anti-FLAG antibody (see below), and two control wells without antibodies. Plates were incubated for 3 h at 4°C and then washed 5 times with 300 μL of ice-cold HNTG, for 10 minutes at 4°C for each wash. After the last wash, 10 μL of PBL buffer (Promega) was added in each well. To measure the input, 2 μl of extract and 8 μL of 1xPBL buffer were put in empty remaining wells. Plates were then incubated 5 minutes at room temperature, and FFL and RL luciferase activities were measured in IP and input wells, using the dual luciferase kit (Promega). Every transfection was performed three times as independent replicates.

To coat the wells of the 96-well plates with M2 anti-FLAG antibodies, High-binding plates were used (Lumitrac, Greiner), and 70 μL of M2 antibody (10 μg/mL in 1xPBS; F1804 Sigma-Aldrich) were put in each well and incubated overnight at room temperature in the dark. The next day, wells were blocked with 300 μL of blocking buffer, for 1 hour at room temperature. IP control wells were treated the same way except that no antibody was put in the well. Blocking buffer was 3% Bovine Serum Albumin (BSA), 5% sucrose, 0.5% Tween 20, 1xPBS). HNTG buffer was 20 mM HEPES-KOH pH 7.9, 150 mM NaCl, 1% Triton X-100, 10% glycerol, 1 mM MgCl2, 1 mM EGTA.

### Co-expression assays and cryo-EM samples preparation

The sequences coding the proteins or protein fragments of interest, wild type and mutants, are subcloned between *Nde*I and *BamH*I rectriction sites into 3 pET-based vectors: pnEA which is ampicillin resistant and encodes a protein carrying an N-terminal 6-Histidine tag followed by a 3C protease site, pnCS which is spectinomycin resistant and encodes a native protein, and pnYK which is kanamycin resistant and encodes also a native protein.

For co-expression assays, the vectors encoding the proteins or protein fragments are co-transformed into *E. coli* BL21(DE3)pRARE2 expression strain. The clones selected on LB-agar plates supplemented with the adequate antibiotics are grown in 100 mL LB medium at 37°C; when OD_600_ reaches 0.7, the cultures are induced with 0.2 mM IPTG and placed O/N at 20°C under agitation.

Then the cells are harvested, and the pellets resuspended into 4 mL lysis buffer (HEPES 25 mM pH 7.5; NaCl 300 mM; imidazole 10 mM; TCEP 0.5 mM), sonicated, and the lysate is cleared by centrifugation. 200 µL TALON resin (50% slurry) are added to the supernatant (soluble fraction) and after a 15 to 30 min binding step on a rotary shaker, the resin is spun down, washed 3 times with 500 µL lysis buffer, and 250 µL elution buffer (HEPES 25 mM pH 7.5; NaCl 300 mM; Imidazole 300 mM; TCEP 0.5 mM) are added to recover the complexes.

Purification steps are followed on SDS-PAGE where insoluble fraction (P), Soluble fraction (So), proteins Bound to the resin (B) and Elution (E) are loaded and colored by Coomassie blue staining.

For reconstituted R2SP and R2S’P cryo-EM samples preparation, R1R2 and S1P2 or S1’P2 are produced in large cultures and purified separately, then mixed.

For R2SP, transformation of pETDuet:His-R1R2-FH8 on one hand, and co-transformation of pnEA::P2 and pnYK::S1 (pnYK::S1_622-926_ for R2S’P) on the other hand are performed into *E. coli* BL21(DE3)pRARE2 expression strain. The clones selected on LB-agar plates supplemented with the adequate antibiotics are grown in 1L flasks in LB medium at 37°C and when OD_600_ reaches 0.7, the cultures are induced with 0.2 mM IPTG. After O/N growth at 20°C under agitation, the cells are harvested and pellets resuspended into lysis buffer (HEPES 25 mM pH 7.5; NaCl 300 mM; Imidazole 10 mM; TCEP 0.5 mM; Glycerol 5% for R1R2; ADP 0.5 mM for R2S’P) (40 mL/L culture pellet), and sonicated. The lysate is cleared by centrifugation and the complexes are purified on TALON resin as followed: 2 mL resin (50% slurry) are added to the 40 mL supernatant (soluble fraction) and after a 15 to 30 min binding step on a rotary shaker, the resin is spun down, washed 3 times with 5 mL lysis buffer and the complexes are eluted with 3 x 5 mL elution buffer (HEPES 25 mM pH 7.5; NaCl 300 mM; Imidazole 300 mM; TCEP 0.5 mM; Glycerol 5% for R1R2; ADP 0.5 mM for R2S’P). Human Rhino Virus 3C (HRV-3C) protease is added to the elution for O/N cleavage at 4°C of the FH8 C-terminal tag for R2 and the Hisx6 N-terminal tag for S1P2 (or S1’P2). After size exclusion chromatography (SEC) on Superose6 column, R1R2 is mixed with excess S1P2 (or S1’P2), and the reconstituted R2SP (or R2S’P) complex is purified again on Superose6 column in final buffer (HEPES 20 mM pH 7.5; NaCl 150 mM; TCEP 0.5 mM; ADP 0.5 mM for R2S’P) to eliminate excess of SP (or S1’P2) and generate a cryo-EM sample at 1.1 mg/mL.

### Protein co-expression and purification of for the SPR experiment

R1 and R2 were produced as described in^15^ with the following modifications: (*i*) R2 has an N-terminal FH8-tag preceded by a HRV-3C cleavage site; (*ii*) Peak fractions from the HisTrap were incubated with 5 mM CaCl_2_ and loaded onto a HiPrepTM Octyl FF 16/10 column (Cytiva). A Superose 6 Increase 10/300 GL equilibrated with 20 mM Tris-HCl pH 8.0, 150 mM NaCl, 5% glycerol, 2 mM MgCl_2_ and 0.5 mM TCEP, was used to separate oligomeric species. R1^D302N^/R2^D299N^ active site mutant variant was purified in the presence of 4 mM ATP, and R1ΔD_II_/R2ΔD_II_ truncated variant complex was purified in the absence of nucleotides.

S1’ carries a C-terminal Flag-tag without a cleavage site, while P2_CS or P2 have a C-terminal StrepTag II preceded by an HRV-3C cleavage site. S1’:P2_CS or S1’:P2 were co-expressed in *E. coli* (DE3*) (Novagen, 71400), with 50 µM IPTG overnight at 18°C in a New Brunswick™ (Innova®) 44R Shaker at 150 rpm. The complexes were immobilized in a 5 mL StrepTactin XT HC (IBA life sciences), previously equilibrated in 20 mM HEPES pH 8, 300 mM NaCl, 0.5 mM TCEP, and eluted with 50 mM Biotin. Peak fractions collected from the StrepTactin XT were injected in a Superdex 200 16/60 XK, allowing the isolation of a heterodimer. Collected fractions from the main peak were diluted to reduce the concentration of NaCl to 50 mM, and further polished in Resource TM Q (Cytiva), and eluted with a linear gradient, allowing the separation of a major peak corresponding to the intact complex (w/o degradation) at approximately 170 mM NaCl. Tag removal from collected peak fractions was performed by incubating 18 h at 4°C with 1% (w/w) HRV-3C protease (Thermo Fisher Scientific, TFS). Digested sample was injected into a 5 mL StrepTactin XT in tandem with 1 mL HisTrap. All purification steps were carried out at room temperature and were monitored by NuPage Bis-Tris gels (Invitrogen, NP0302).

### Surface Plasmon resonance

S1’P2_CS, or S1’P2 protein constructs were immobilized onto CM5 (Series S) sensor chips using standard amine coupling. 10 mM HEPES, pH 7.4, 150 mM NaCl (HSB-N) was used as the background buffer during immobilization. The carboxymethyl dextran surface of all flow cells was activated with 20 mM EDC and 5 mM NHS for 1.5 min. Both S1/P2 constructs were injected on the chip surface at a final concentration of 0.5 µg/mL in 10 mM Sodium Acetate buffer pH 5.5. The enzymes were coupled to the surface with a 1 to 2 min injection time at a flow rate of 10 μL min^-1^. The remaining activated groups were blocked with a 5-min injection of 1.0 M ethanolamine, pH 8.5. Typically, 100 ± 10 response units (RU) were obtained for the immobilization of all protein constructs. Negative controls were performed by immobilizing either BSA (TFS) or human Cyclophilin D_43-207_ (CypD) with the same RU levels as the proteins of interest. CypD_43-207_ is a 22 kDa in-house purified protein with similar size to S1_C and confirmed to be active through binding to Cyclosporin A. All R1-R2 complexes were directly dissolved in running buffer (20 mM NaKPi pH 7.5, 150 mM NaCl, 5 mM MgCl_2_, 1 mM DTT, 0.05% P20), in the absence or presence of nucleotides and analyzed with a Biacore 4000 (Biacore AB, GE Healthcare Life Sciences, Uppsala, Sweden). RUVBL complex was 300 s sample injection (30 µL min^-1^) followed by 600 s of buffer flow (dissociation phase). All sensorgrams were processed by subtracting the binding response recorded from the control surface (reference spot), followed by subtracting of the blank buffer injection from the reaction spot. All datasets fit to a simple 1:1 interaction model to determine kinetic (*k*_a_, *k*_d_, K_D_) rate constants and affinity at steady state (K_Dss_).

### ATPase assay

The steady-state ATPase activity of R1R2 complex was evaluated using a coupled enzymatic assay (PK/LDH) that monitors the oxidation of NADH, as described previously (7), with a modified reaction buffer (50 mM HEPES/KOH pH=7.5, 100 mM KCl, 5 mM ATP; 5 mM MgCl_2_) in 96-well plates on a CLARIOstar reader (BMG Labtech) at 37°C. The R1R2 concentration was 1 µM of hexamer. The apparent affinity constant between R1R2 and R3P2 or S1P2 was calculated by relative specific ATPase activity (SA) of R1R2 and fitted with the following equation using R software ^41^: 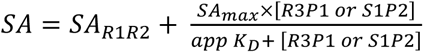

### Fluorescence polarization assay

The fluorescence polarization of BODIPY® FL ATPγS (Jena Bioscience) was measured by Clariostar (filters: excitation: 482-16; dichroic: LP 504; emission: 530-40) on 384 wells-plate at 25°C. All fluorescence polarization assays were performed into FP buffer (25 mM HEPES/KOH pH=7.5, 150 mM KCl, 5 mM MgCl_2_) with a final concentration of BDP-ATPγS at 10 nM. The nucleotides of R1R2 complex were removed by incubation with Agarose-phosphatase alkaline (P0762, Sigma) at 25°C overnight. For binding and equilibrium assays, the proteins were deposited into the wells using a pipette, and the BDP-ATPγS were added via the injector of Clariostar. For exit assays, proteins and BDP-ATPγS were deposited together with a pipette on the wells and unlabeled nucleotides were added via the injector of Clariostar. The data were fitted using R with these equations: (binding assay) 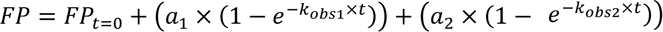; (exit assay) 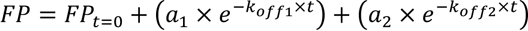

### Native Mass Spectrometry

Samples were buffer exchanged against ammonium acetate (200 mM, pH 7.5) with Zeba Spin desalting columns (7 kDa cutoff, TFS, Rockford, IL, USA). Sample concentrations were determined by UV absorbance using a NanoDrop 2000 spectrophotometer (TFS, France). Prior to injection, full length R1R2 and S’P2 were diluted, and mixed in ratio 1:2 (detected species were the same even with higher proportions of S1’P2). nMS experiments were carried out on a hybrid electrospray quadrupole time-of-flight mass spectrometer (Synapt G2 HDMS, Waters, Manchester, UK) coupled to an automated chip-based nanoelectrospray source (Triversa Nanomate, Advion, Ithaca, USA) operating in the positive ion mode. Mass spectrometer calibration was performed from 1,000 to 20,000 m/z using singly charged ions produced by a 2 mg/mL solution of cesium iodide in 2-propanol/water (v/v). Instrumental parameters were optimized to ensure the transmission of high masses while maintaining non-covalent interactions. The backing pressure was set to 6 mbar and the cone voltage to 200 V. Data interpretation was performed using MassLynx v4.1 (Waters, Manchester, UK).

### Mass Photometry

Measurements of were done on a TWO MP (Refeyn Ltd, Oxford, UK) at room temperature. We used microscope slides (24 x 50 mm, 170 ± 5 µm, No. 1.5H, Paul Marienfeld GmbH & Co. KG, Germany) that were cleaned (milli-Q water, isopropanol, milli-Q water) and dried (clean nitrogen stream) and further assembled with six-well silicone gaskets (cut from CultureWellTM, 50-3 mm DIA x 1 mm Depth, 3-10 µL, Grace Bio-Labs, Inc., Oregon, USA). Focus was made each time with one 18 µL PBS droplet before mixing with 2 µL of sample to analyze. A contrast-to-mass calibration was performed twice a day with a mix of BSA, Bevacizumab and Glutamate Dehydrogenase in PBS. For R1R2 and R2SP analysis, samples were first diluted in their native sample to reach 500 nM and 2 µL of these stock solutions were then quickly diluted to 50 nM in the 18 µL droplet of PBS. Three movies of 60 s (3 000 frames) were recorded for each sample using AcquireMP sotware (Refeyn Ltd, Oxford, UK), data were processed with DiscoverMP software (Refeyn Ltd, Oxford, UK).

### XL-MS

SP heterodimer and R2SP complex (20 mM HEPES; 150 mM NaCl; 0.5 mM TCEP; pH 7.5) were both incubated in biological triplicate with DSBU cross-linker (CF plus Chemicals) at 100 molar excesses for 45 min (RT) and quenched with 20 mM ammonium bicarbonate for 20 min. XL reaction was controlled using home-made SDS-PAGE (12%). Samples were then reduced for 30 min at 37°C (5 mM DTT), alkylated for 1 h in the dark (15 mM Iodoacetamide), and finally digested overnight at 37°C with Trypsin/Lys-C(Promega, Madison, USA) at a 50:1 Protein:Enzyme ratio. Digestion was quenched with 1% v/v TriFluoroacetic Acid (TFA), and peptides were submitted to a SPE cleanup (Sep-Pak C18 1cc, 50 mg). After evaporation and dilution in 2% ACN/0.1% (Formic Acid) FA, samples were analyzed in nanoLC-MS/MS using a nanoAcquity UPLC (Waters, Milford, USA) coupled to a Q-Exactive HF-X Orbitrap mass spectrometer (TFS) controlled by XCalibur software (v4.0.27.9). Trapping was done on a nanoEase M/Z Symmetry C18 precolumn (180 µm x 20 mm, 5 µm particle size, Waters) and separation on a nanoEase M/Z Peptides BEH C18 (75 µm x 250 mm, 1.7 µm particle size, Waters). The following 105 min gradient was applied using mobile phase A (water/0.1 % FA) and B (ACN/0.1% FA): 3% for 5 min, 3-40% B in 90 min, 40-99% B in 2 min, 99% B for 5 min, 99-3% B in 2 min. Flow rate was set to 400 nL/min and column temperature to 60°C. Full scan MS spectra were acquired in positive mode with a resolution of 120 000, with a 300-1 800 m/z range. Fragmentation of Top 10 most intense ions (charge states 3-8 +) were done using HCD stepped collision energy (27, 30, 33% NCE). MS/MS spectra were acquired with a resolution of 30 000 and a 2 m/z isolation window. Raw data were converted into .mgf files and processed using MeroX software v2.0.1.4 with a mass tolerance of 5 ppm for precursor ions and 10 ppm for product ions. A 5% FDR cut-off and a signal-to-noise of at least 2 were also applied. Proteases cleavage sites were defined as Lys and Arg with maximum 3 missed cleavages, only peptides of at least 5 residues were considered. Cysteine carbamidomethylation was set as fixed modification and Methionine Oxidation as variable modification (2 max. mod.). Lys and N-ter as well as Ser, Thr and Tyr were considered as cross-linking sites. Database containing interest proteins and > 50 common contaminants was used for identification with RISEUP mode and 1 max. missing ion. Cross-links composed of consecutive amino acid sequences were not used and all cross-links identified were manually validated. Only cross-links identified in at least 2/3 XL replicates were validated. The SP and R2SP XL-MS datasets have been deposited to ProteomeExchange via the PRIDE partner repository, with dataset identifier PXD056494. For mapping on sequences and distance validation of identified XLs on the different structures (< 32 Å cutoff for Cα-Cα), we used the xiVIEW webserver (https://xiview.org)^42^. XLs were further visualized using PyMol version 2.5.4 (Schrödinger, LLC)^40^ and UCSF ChimeraX version 1.8^43^.

### Grid preparation and cryo-EM images acquisition

Cryo-EM grids were prepared and systematically checked at METI, Toulouse. Immediately after glow discharge, 3.5 µL of purified R2S’P particles (at 0.45 mg/mL as estimated by Bradford) were deposited onto QUANTIFOIL holey carbon grids (R2/1, 300 Mesh). Grids were plunge-frozen using a Leica EM-GP automat; the temperature and humidity level of the loading chamber were maintained at 20°C and 95%, respectively. Excess solution was blotted with a Whatman filter paper no. 1 for 1.7–1.9 s, and grids were immediately plunged into liquid ethane (**-**183°C).

Images were recorded on a Titan Krios electron microscope (FEI, TFS) located at EMBL, Heidelberg, Germany. The cryo-electron microscope operated at 300 kV and was equipped with a Gatan K2 summit direct electron detector using counting mode. Automatic image acquisition was performed with SerialEM, at a magnification corresponding to a calibrated pixel size of 0.645 Å and a total electron dose of 51.92 e-/Å2 over 40 frames. Nominal defocus values ranged from **-**0.8 μm to **-**2.8 μm.

### Single-particle analysis

Image analysis was performed using Relion 3.1^44^, unless otherwise stated. An overview of the image processing scheme is presented in figures S8 and S11. Sixteen thousand five hundred and sixty-six stacks of frames were collected at EMBL Heidelberg. Frame stacks were aligned to correct for beam-induced motion using MotionCor2^45^. Contrast transfer function (CTF) and defocus estimation were performed on the realigned stacks using GCTF^46^. After selection upon CTF estimation quality, maximum resolution on their power spectra, and visual checking, ‘good’ micrographs were retained for further analysis. A total of 2,158,039 particles were initially picked and then extracted in boxes of 304 × 304 pixels, using the LogPicker Autopick option from Relion 3.1^44^. A first 2D classification was performed (on particle images binned by a factor of 4) to sort out bad particles, yielding a total of 1,870,648 remaining particles. Of note, on the 1,870,648 picked particles, at least 1,303,376 were classified as top views; these preferential orientations thus represent at least 70% of the imaged particles, suggesting a strong air-water interface bias during plunge freezing. A subset of 2D classes were first used for an *ab initio* 3D reconstruction, and the resulting dodecameric 3D map was used as a reference for a 3D classification in five classes of the particle dataset. One class harbored a dodecameric form of the R2SP with a good level of detail. The 253,429 particles from this class were re-extracted without imposing any binning factor, and a consensus 3D structure was obtained using RELION’s 3D auto-refine option, that reached a resolution of 3.5 Å according to gold-standard Fourier shell correlation (FSC) at 0.143 after CTF Refinement and particle polishing. A post-processed map of the dodecameric assembly was calculated using DeepEMhancer^45^. The remaining classes were merged and subjected to a 3D classification (K=5), using an hexameric R1R2 atomic model derived from (PDB-ID: 2XSZ)^24^ as reference. One of the five 3D classes resulted in a clearly defined hexameric shape typical to the R1R2 assembly. The corresponding 302,119 particles were re-extracted without imposing any binning factor, and a consensus 3D structure was obtained at 3.6 Å resolution according to gold-standard FSC procedure. To try to sort out structural heterogeneity within these hexameric particles, two different strategies were used (fig. S11). The first was to perform a global, unmasked 3D classification (K=5) while skipping orientation searches, and the second used a mask around a monomer of R1+R2+S1_C for a masked and focused 3D classification while skipping orientation searches. From the first strategy, a “cleaned” 3D class of 167,033 particles was obtained. These particles were subjected to 3D auto refinement, and the maps were subsequently improved by particle-based motion correction, B-Factor weighting, and CTF refinement, resulting in a final resolution of 3.6 Å that displayed a hexameric ring crowned by at least one “horn” (corresponding to S1_C).

To assess how many S1_C could be associated with R1R2 hexameric ring, the second strategy (right panel on fig. S11) was to apply a mask around a monomer of R1, R2, S1_C, and perform a focused 3D classification while skipping orientation searches. This resulted in 5 classes with either one or two horn-like structures of S1_C which were refined to resolutions ranging from 5.0 to 3.8 Å. Local resolution of the dodecameric and hexameric assemblies presented in figures S8 and S11 were assessed using ResMap^47^.

### Structural heterogeneity assessment within the cryo-EM datasets

To characterize their nature and level of structural heterogeneity, 302,119 particles corresponding to hexameric, or 252,248 particles corresponding to dodecameric assemblies (dodecamer before polishing) were also subjected to AI-based sorting by cryoDRGN version 2.3.0^34^. All models were trained for 50 epocs, parallelized through several GPU’s. Both hexameric and dodecameric particles were downsampled to a box size of 128 pixels with a pixel size of 1.52 Å/pixel for evaluation. We trained a cryoDRGN model on these downsampled particles with three layers and 256 nodes per layer for both the encoder and decoder network on layers with 512 nodes). This resulted in a smooth distribution of Z values, indicating that heterogeneity in the dataset originates purely from continuous conformational changes and not from populations of particles with distinct compositions, in agreement with the 3D classification results using Relion. We then represented the maps (movies S1 and S2) using the traversal of the first, second, and third principal components of variability in the Z values, as implemented in cryoDRGN. For the hexameric particles, the resulting maps show a variety of states, including states with different number of S1_C’s decorating the “top” region of the ring, the presence of density “beneath” the R1R2 ring that might correspond to P2, D_II_ from R1 moving laterally, and lack of density for D_II_ of R2 that are bound to S1_C. For the dodecameric particles, the resulting maps show different ways that both hexamers might move relative to each other, including twisting, compaction, and tilting, but also D_II_ density variations, specifically from R2.

### Cryo-EM Maps Interpretation

Atomic models of the R1R2 hexamer (PDB-ID 6H7X, https://doi.org/10.2210/pdb6H7X/pdb; 2C9O, https://doi.org/10.2210/pdb2C9O/pdb; and 6FO1, https://doi.org/10.2210/pdb6FO1/pdb) as well as of the NMR structure of S1_C (this work) herein presented were first fitted in the cryo-EM maps of interest as rigid body using the “fit” command in UCSF Chimera^48^. Manual refinements and adjustments, as well as flexible and jiggle fittings were then realized on various chains of the models in Coot^49^. Final atomic model was refined using REFMAC5^50^. Final model evaluation was done with MolProbity^51^. Overfitting statistics were calculated by a random displacement of atoms in the model, followed by a refinement against one of the half-maps in REFMAC5, and Fourier shell correlation curves were calculated between the volume from the atomic model and each of the half-maps in REFMAC5 (table S5) Maps and models visualization was done with Coot, UCSF Chimera and ChimeraX^43^. Figures and movies were created using ChimeraX and Kdenlive (https://kdenlive.org/en/).

### R2S’P integrative model establishment

The 3D structures of P2 and S1, either in their full-length form or cut into smaller domains, were predicted using AlphaFold3^52^. Namely, the following aminoacid sequences of S1’ (622-926) and S1’_TPR3_plus (622-780); or P2 (1-315), P2_PIH_plus (1-163), P2_CS_plus (160-315) were submitted to structure prediction by AF3 (Fig. 1). AF3 structural prediction of P2’s PIH domain was fitted on a segmented low-resolution cryo-EM map presented in movie S2 using Chimera’s Segger option. Then, combinations of S1’:P2 (full length or fragments above described) were systematically submitted to AF3. Finally, different sequences enclosing R1 and R2 D_IIs_ (namely: R1_DII_: 120-296; R1_DIIext_: 120-240; R2_DII_: 126-295; R2_DIIext_: 126-232) were also added to S1’:P2 complexes, yielding S1’:P2:R1R2 assemblies for AF3 prediction. Assays using different copy numbers of these proteins/domains were also systematically performed (S1’ and P2: 1 or 2 copies; R1 and R2: 1, 2 or 3 copies of each protein segment). For each of these AF3 predictions, all 5 output structures were examined using Chimera. The final structures of P2’s CS and S1’ TPR3 domains that were retained for building the integrative R2S’P model corresponded to the 4^th^ best AF3 prediction of the following construct, 2xS1’_TPR3_plus:2xP2:1xR1_DIIext_:1xR2_DIIext_, because it was in best agreement with the experimental data from cryo-EM, NMR and XL-MS. Atomic coordinates of the P2’s CS and S1’ TPR3 domains were extracted from this AF3 prediction and added to the R2S cryo-EM atomic model supplemented with P2’s PIH domain using Coot and Chimera.

## Acknowledgments

The authors would like to thank S. Balor and V. Soldan (METI platform, Toulouse, member of the national infrastructure France-BioImaging supported by the French National Research Agency (ANR-10-INBS-04)), for technical support in electron microscopy, and the engineers from the CALMIP HPC facility. We thank the Biophysics and Structural Biology (B2S) platform of SMP (“Service Mutualisé de Plateformes, Université de Lorraine”, https://umsibslor.univ-lorraine.fr/en/facility/biophysics-structural-biology-b2s) for access to the NMR and SLS core facilities.

## Author contributions

P.E.S., M.E.C., M.L., H.G.F., T.C., E.D., L.P., A.C.F.P. performed the experiments. P.E.S., M.E.C., H.G.F., P.M.F.S., C.V., T.M.B., P.M., S.C., X.M., C.P.C., M.Q. conceived the study. P.M.F.S., C.V., T.M.B., P.M., S.C., X.M., C.P.C., M.Q. supervised the work. P.E.S., M.E.C., H.G.F., A.C.F.P., P.M.F.S., C.V., T.M.B., P.M., S.C., X.M., C.P.C., M.Q. wrote the manuscript. All co-authors proofread the manuscript.

## Data and materials availability

All data needed are present in the paper and/or the Supplementary Materials. Additional data related to this paper may be requested from the authors. The cryo-EM maps and the models have been deposited in the Electron Microscopy Data Bank (EMDB) with accession codes EMD-52013 and EMD-52336 for the hexameric R2S’P and dodecameric R1R2 cryo-EM maps, respectively, and PDB ID 9HB4 and 9HP0 for the R1R2S1_C and dodecameric R1R2 models, respectively. The chemical shifts and NMR structure of the human S1_C are deposited in BMRB (ID: 34968) and PDB (ID: 9HKR) respectively. The mass spectrometry proteomics data have been deposited to the ProteomeXchange Consortium via the PRIDE^53^ partner repository with the dataset identifier PXD056494.

## Figures and Tables

**Table S1.**
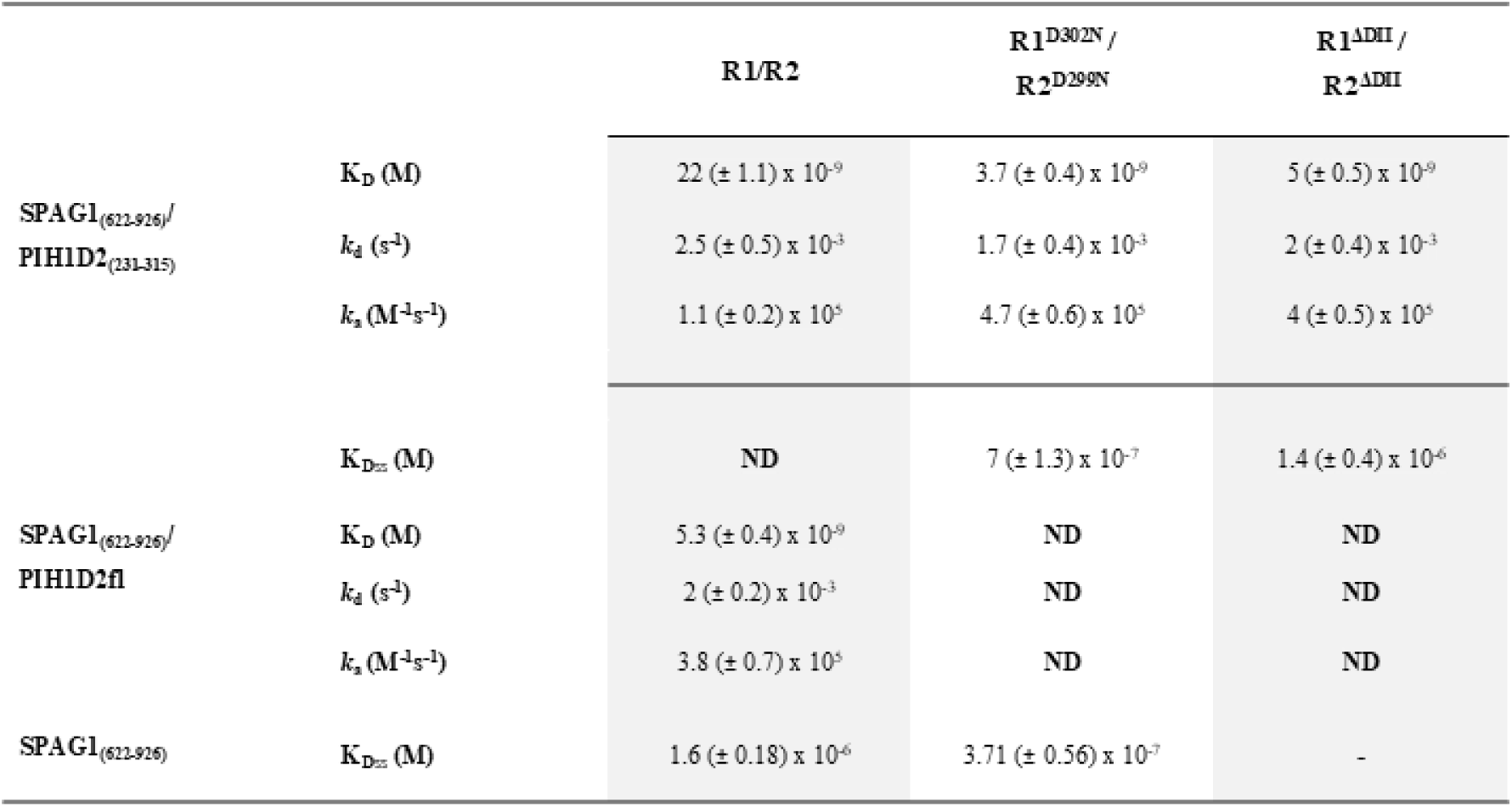
Affinity (K_Dss_) and Kinetic (*k*_a_, *k*_d_, K_D_) interaction parameters as determined by SPR. ΔD_II_ means R1 or R2 without the D_II_.

**Table S2.** NMR and refinement statistics for top-20 S1_C structures.

| NMR distance and dihedral constraints |  | S1_C |
| --- | --- | --- |
| Distance constraints |  |  |
| Total NOE |  | 3504 |
| Intra-residue |  | 892 |
| Inter-residue |  | 2612 |
| Sequential ($\vert i - j\vert = 1$) | | 818 |
| Medium-range ($\vert i - j\vert \leq 4$) | | 947 |
| Long-range ($\vert i - j\vert \geq 5$) | | 847 |
| Total dihedral angle restraints |  | 232 |
| $\phi$ | | 113 |
| $\psi$ | | 119 |
| Structure statistics after AMBER refinement |  |  |
| Violations |  |  |
| Distance constraints (Å) |  | 0.11 ± 0.01 |
| Dihedral angle constraints (°) |  | 4.04 ± 1.99 |
| Max. dihedral angle violation (°) |  | 6.36 |
| Max. distance constraint violation (Å) |  | 0.12 |
| Violation occurrences |  |  |
| Distances constraints (> 0.2 Å) |  | 0 |
| Dihedral angle constraints (> 5°) |  | 1.85 ± 1.39 |
| R.m.s. deviations from idealized geometry |  |  |
| Bond lengths (Å) |  | 0.013 ± 0.00 |
| Bond angles (°) |  | 1.90 ± 0.05 |
| R.m.s. deviation to best structurea (Å) |  |  |
| All heavy atoms |  | 1.19 ± 0.12 |
| All backbone atoms |  | 1.65 ± 0.14 |
| Heavy atoms in secondary structures |  | 0.88 ± 0.06 |
| Backbone atoms in secondary structures |  | 0.39 ± 0.06 |

**Table S3.**
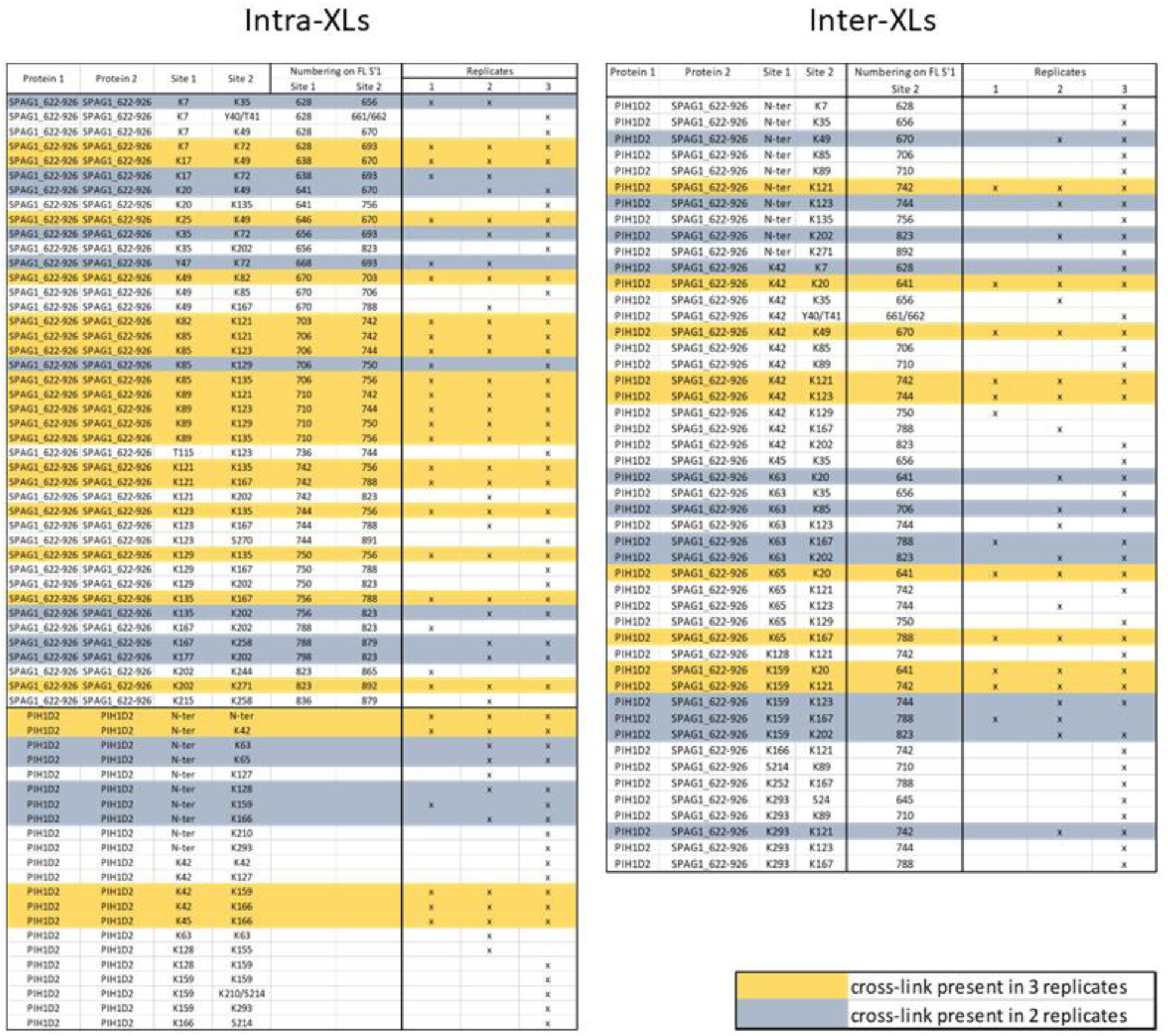
List of identified S1’P2 intra- and inter protein cross-links identified. Validated XL-pairs present in 2/3 and 3/3 replicates are highlighted respectively in yellow and grey.

**Table S4.**
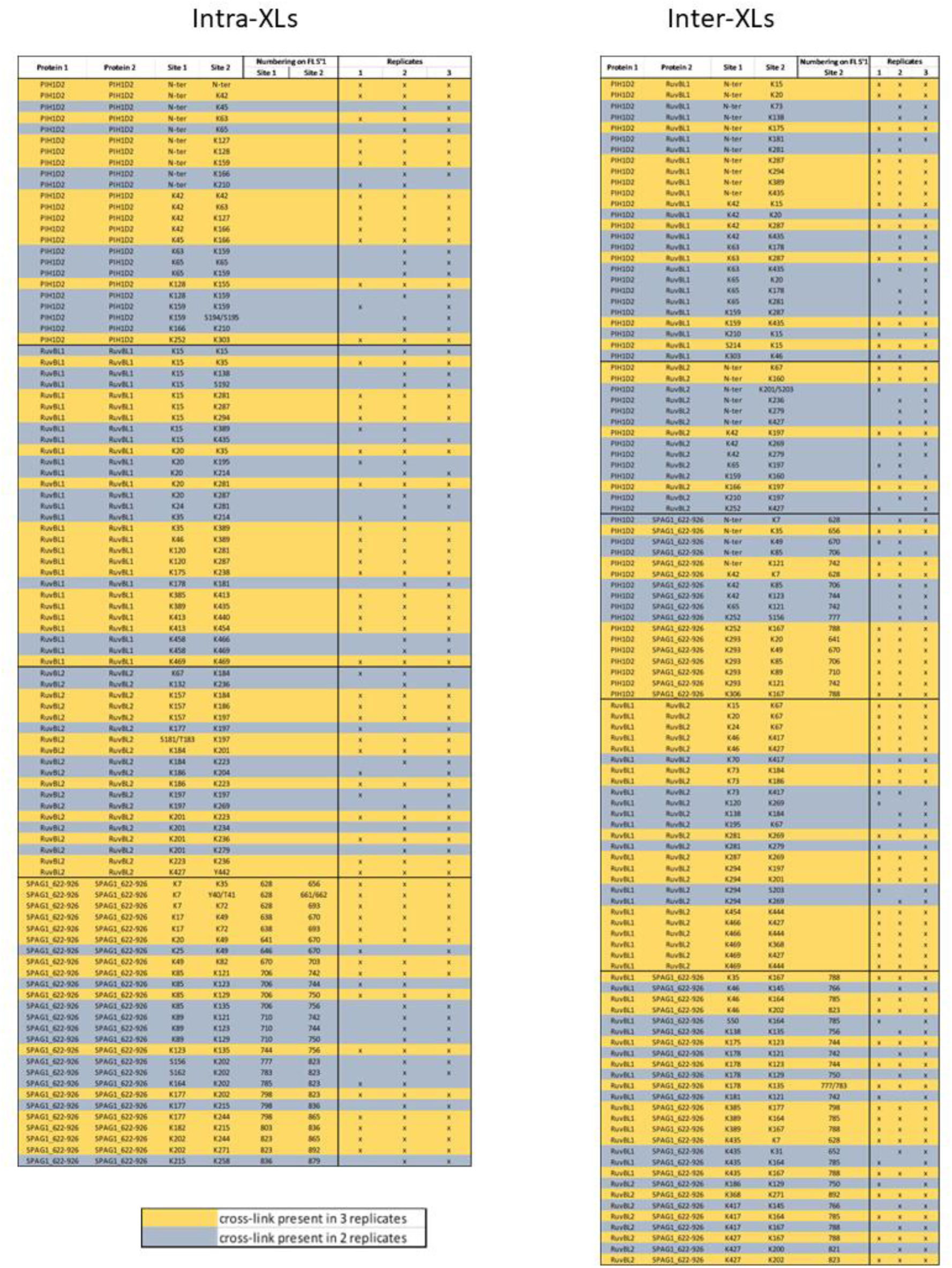
List of unique XLs identified in replicates of R2S’P XL-MS experiments. Validated XL-pairs present in 2/3 and 3/3 replicates are highlighted respectively in yellow and grey.

**Table S5.** Cryo-EM data collection, atomic model refinement and validation statistics.

|  | Hexameric R1R2 + SPAG1_C | Dodecameric R1R2 |
| --- | --- | --- |
| Microscope model | FEI Titan Krios cryo-transmission electron microscope |  |
| Detector model | Gatan K2 summit direct electron detector |  |
| Number of datasets | 1 |  |
| Number of micrographs collected | 16,566 |  |
| Pixel size (Å) | 0.645 |  |
| Defocus range (µm) | 0.8 – 2.8 |  |
| Voltage (kV) | 300 |  |
| Electron dose (e<sup>-</sup> Å<sup>-2</sup>) | 51.92 |  |
| Name of 3D reconstruction/ model | R1R2_S1_C | Dodecameric R1R2 |
| EMDB entry of map | EMD-52013 | EMD-52336 |
| PDB entry of the full model | 9HB4 | 9HPO |
| Final number of particles | 302,119 | 235,429 |
| Resolution (Å) (FSC threshold = 0.143) | 3.6 | 3.5 |
| Map sharpening B-factor (Å<sup>2</sup>) | -126 | -152 |
| **Refinement and model validation statistics<sup>(a)</sup>** |  |  |
| Model refinement resolution range (Å) | 40-3.0 | 40-2.8 |
| Model resolution (Å) (FSC threshold = 0.5) | 3.68 | 3.8 |
| Clashscore (all atoms) | 9.34 | 47.88 |
| Model Composition |  |  |
| Non-hydrogen atoms | 19008 | 37665 |
| Protein residues | 2430 | 4783 |
| Ligands | 6 ADP | 12 ADP |
| MolProbity Score | 2.23 | 2.76 |
| Protein |  |  |
| Rmsd (bonds lengths, Å) | 0.003 | 0.003 |
| Rmsd (angles, °) | 0.728 | 0.680 |
| Ramachandran plot (%) |  |  |
| favored | 94.82 | 95.01 |
| allowed | 5.14 | 4.86 |
| outliers | 0.04 | 0.13 |
| <sup>(a)</sup>Models were validated using MolProbity implemented in PHENIX.REFINE (Adams et al., 2010) |  |  |

## REFERENCES

1. Houry, W. A., Bertrand, E. & Coulombe, B. The PAQosome, an R2TP-Based Chaperone for Quaternary Structure Formation. Trends in Biochemical Sciences 43, 4–9 (2018).

2. Boulon, S. et al. HSP90 and its R2TP/Prefoldin-like cochaperone are involved in the cytoplasmic assembly of RNA polymerase II. Mol Cell 39, 912–24 (2010).

3. Cloutier, P. et al. R2TP/Prefoldin-like component RUVBL1/RUVBL2 directly interacts with ZNHIT2 to regulate assembly of U5 small nuclear ribonucleoprotein. Nature Communications 8, 15615 (2017).

4. Coulombe, B., Cloutier, P. & Gauthier, M. S. How do our cells build their protein interactome? Nature communications 9, 2955 (2018).

5. Cloutier, P. et al. Upstream ORF-Encoded ASDURF Is a Novel Prefoldin-like Subunit of the PAQosome. J. Proteome Res. 19, 18–27 (2020).

6. Ewens, C. A. et al. Architecture and Nucleotide-Dependent Conformational Changes of the Rvb1-Rvb2 AAA+ Complex Revealed by Cryoelectron Microscopy. Structure 24, 657–66 (2016).

7. Henri, J. et al. Deep Structural Analysis of RPAP3 and PIH1D1, Two Components of the HSP90 Co- chaperone R2TP Complex. Structure (2018) doi:10.1016/j.str.2018.06.002.

8. Horejsi, Z. et al. CK2 phospho-dependent binding of R2TP complex to TEL2 is essential for mTOR and SMG1 stability. Mol Cell 39, 839–50 (2010).

9. Boulon, S. et al. The Hsp90 chaperone controls the biogenesis of L7Ae RNPs through conserved machinery. J Cell Biol 180, 579–95 (2008).

10. Machado-Pinilla, R., Liger, D., Leulliot, N. & Meier, U. T. Mechanism of the AAA+ ATPases pontin and reptin in the biogenesis of H/ACA RNPs. Rna 18, 1833–45 (2012).

11. Schlotter, F. et al. Proteomic analyses reveal new features of the box H/ACA RNP biogenesis. Nucleic Acids Research gkad129 (2023) doi:10.1093/nar/gkad129.

12. Bizarro, J. et al. NUFIP and the HSP90/R2TP chaperone bind the SMN complex and facilitate assembly of U4- specific proteins. Nucleic acids research 43, 8973–89 (2015).

13. Seraphim, T. V. et al. Assembly principles of the human R2TP chaperone complex reveal the presence of R2T and R2P complexes. Structure 30, 156–171.e12 (2022).

14. Palacios-Abella, A. et al. The structure of the R2T complex reveals a different architecture of the related HSP90 co-chaperones R2T and R2TP. 2024.03.27.587014 Preprint at 10.1101/2024.03.27.587014 (2024).

15. Maurizy, C. et al. The RPAP3-Cterminal domain identifies R2TP-like quaternary chaperones. Nature communications 9, 2093 (2018).

16. Rodríguez, C. F. & Llorca, O. RPAP3 C-Terminal Domain: A Conserved Domain for the Assembly of R2TP Co-Chaperone Complexes. Cells 9, 1139 (2020).

17. Chagot, M. E. et al. Binding properties of the quaternary assembly protein SPAG1. The Biochemical journal 476, 1679–1694 (2019).

18. Dermouche, S., Chagot, M.-E., Manival, X. & Quinternet, M. Optimizing the First TPR Domain of the Human SPAG1 Protein Provides Insight into the HSP70 and HSP90 Binding Properties. Biochemistry (2021) doi:10.1021/acs.biochem.1c00052.

19. Dos Santos Morais, R., et al. Deciphering cellular and molecular determinants of human DPCD protein in complex with RUVBL1/RUVBL2 AAA-ATPases. Journal of Molecular Biology 434, 167760 (2022).

20. Munoz-Hernandez, H. et al. Structural mechanism for regulation of the AAA-ATPases RUVBL1-RUVBL2 in the R2TP co-chaperone revealed by cryo-EM. Science advances 5, eaaw1616 (2019).

21. Ayala, R. et al. Structure and regulation of the human INO80–nucleosome complex. Nature 556, 391–395 (2018).

22. Silva, S. T. N. et al. X-ray structure of full-length human RuvB-Like 2 – mechanistic insights into coupling between ATP binding and mechanical action. Sci Rep 8, 13726 (2018).

23. Theobald, D. L., Mitton-Fry, R. M. & Wuttke, D. S. Nucleic acid recognition by OB-fold proteins. Annu Rev Biophys Biomol Struct 32, 115–133 (2003).

24. Gorynia, S. et al. Structural and functional insights into a dodecameric molecular machine - the RuvBL1/RuvBL2 complex. Journal of structural biology 176, 279–91 (2011).

25. López-Perrote, A. et al. Regulation of RUVBL1-RUVBL2 AAA-ATPases by the nonsense-mediated mRNA decay factor DHX34, as evidenced by Cryo-EM. eLife 9, e63042 (2020).

26. Yamamoto, R., Hirono, M. & Kamiya, R. Discrete PIH proteins function in the cytoplasmic preassembly of different subsets of axonemal dyneins. The Journal of cell biology 190, 65–71 (2010).

27. Knowles, M. R. et al. Mutations in SPAG1 cause primary ciliary dyskinesia associated with defective outer and inner dynein arms. American journal of human genetics 93, 711–20 (2013).

28. Li, Y., Zhao, L., Yuan, S., Zhang, J. & Sun, Z. Axonemal dynein assembly requires the R2TP complex component Pontin. Development 144, 4684–4693 (2017).

29. Martino, F. et al. RPAP3 provides a flexible scaffold for coupling HSP90 to the human R2TP co-chaperone complex. Nature communications 9, 1501 (2018).

30. Tian, S. et al. Pih1p-Tah1p Puts a Lid on Hexameric AAA+ ATPases Rvb1/2p. Structure 25, 1519–1529 e4 (2017).

31. Rivera-Calzada, A. et al. The Structure of the R2TP Complex Defines a Platform for Recruiting Diverse Client Proteins to the HSP90 Molecular Chaperone System. Structure (2017) doi:10.1016/j.str.2017.05.016.

32. Paiva, A. C. F. et al. Extract2Chip—Bypassing Protein Purification in Drug Discovery Using Surface Plasmon Resonance. Biosensors 13, 913 (2023).

33. Scheres, S. H. W. A Bayesian view on cryo-EM structure determination. J Mol Biol 415, 406–418 (2012).

34. Zhong, E. D., Bepler, T., Berger, B. & Davis, J. H. CryoDRGN: reconstruction of heterogeneous cryo-EM structures using neural networks. Nat Methods 18, 176–185 (2021).

35. Lopez-Perrote, A., Munoz-Hernandez, H., Gil, D. & Llorca, O. Conformational transitions regulate the exposure of a DNA-binding domain in the RuvBL1-RuvBL2 complex. Nucleic acids research 40, 11086–99 (2012).

36. Smith, A. J. et al. The role of SPAG1 in the assembly of axonemal dyneins in human airway epithelia. Journal of Cell Science 135, jcs259512 (2022).

37. Yamaguchi, H., Oda, T., Kikkawa, M. & Takeda, H. Systematic studies of all PIH proteins in zebrafish reveal their distinct roles in axonemal dynein assembly. eLife 7, e36979 (2018).

38. Yenerall, P. et al. RUVBL1/RUVBL2 ATPase Activity Drives PAQosome Maturation, DNA Replication and Radioresistance in Lung Cancer. Cell Chemical Biology 27, 105–121.e14 (2020).

39. Horejsi, Z. et al. Phosphorylation-dependent PIH1D1 interactions define substrate specificity of the R2TP cochaperone complex. Cell reports 7, 19–26 (2014).

40. Schrodinger, L. The PyMOL Molecular Graphics System, Version 1.8. (2015).

41. 41. R Core Team (2020). European Environment Agency https://www.eea.europa.eu/data-and-maps/indicators/oxygen-consuming-substances-in-rivers/r-development-core-team-2006.

42. Combe, C. W., Graham, M., Kolbowski, L., Fischer, L. & Rappsilber, J. xiVIEW: Visualisation of Crosslinking Mass Spectrometry Data. Journal of Molecular Biology 436, 168656 (2024).

43. Goddard, T. D. et al. UCSF ChimeraX: Meeting modern challenges in visualization and analysis. Protein Science 27, 14–25 (2018).

44. Zivanov, J. et al. New tools for automated high-resolution cryo-EM structure determination in RELION-3. eLife 7, e42166 (2018).

45. Zheng, S. Q. et al. MotionCor2: anisotropic correction of beam-induced motion for improved cryo-electron microscopy. Nat Methods 14, 331–332 (2017).

46. Zhang, K. Gctf: Real-time CTF determination and correction. Journal of Structural Biology 193, 1–12 (2016).

47. Kucukelbir, A., Sigworth, F. J. & Tagare, H. D. The Local Resolution of Cryo-EM Density Maps. Nat Methods 11, 63–65 (2014).

48. Pettersen, E. F. et al. UCSF Chimera—A visualization system for exploratory research and analysis. Journal of Computational Chemistry 25, 1605–1612 (2004).

49. Brown, A. et al. Tools for macromolecular model building and refinement into electron cryo-microscopy reconstructions. Acta Cryst D 71, 136–153 (2015).

50. Murshudov, G. N. et al. REFMAC5 for the refinement of macromolecular crystal structures. Acta Cryst D 67,

51. Chen, V. B. et al. MolProbity: all-atom structure validation for macromolecular crystallography. Acta Cryst D 66, 12–21 (2010).

52. Abramson, J. et al. Accurate structure prediction of biomolecular interactions with AlphaFold 3. Nature 630, 493–500 (2024).

53. Perez-Riverol, Y. et al. The PRIDE database resources in 2022: a hub for mass spectrometry-based proteomics evidences. Nucleic Acids Research 50, D543–D552 (2022).

